# Integrative Analysis of Neuroimaging and Microbiome Data Predicts Cognitive Decline in Parkinson’s Disease

**DOI:** 10.1101/2025.03.28.645180

**Authors:** Büşranur Delice, Özkan Ufuk Nalbantoğlu, Süleyman Yıldırım

## Abstract

Parkinson’s disease (PD) is a neurodegenerative disorder characterized by motor and non-motor symptoms, including cognitive impairment (CI) ranging from mild cognitive impairment (MCI) to Parkinson’s disease dementia (PDD). Growing evidence supports the gut-brain axis as playing an essential role in the pathophysiology of PD, suggesting promising applications for combining advanced neuroimaging techniques with microbiome profiling to accelerate biomarker discovery and therapeutic innovation. This study combines resting-state functional magnetic resonance imaging (rs-fMRI) and 16S rRNA sequencing of stool and saliva to identify biomarkers predictive of CI in PD.

A stepwise feature selection pipeline, incorporating ANOVA, random forest ranking, and partial correlation analysis, was applied to extract biologically meaningful features from rs-fMRI connectivity matrices and microbial taxa. Independent and joint machine learning models, including Random Forest, support vector machine, XGBoost, and logistic regression, were evaluated for their predictive performance. The joint model, integrating neuroimaging and microbiome features, outperformed modality-specific models in classifying HC, MCI, and PDD stages, achieving an accuracy of 88.9% and AUC of 97.2% with Random Forest. Key fMRI features involved the salience and default mode networks, while microbial biomarkers included taxa such as *Faecalibacterium, Veillonella,* and *Streptococcus.* Correlations between microbial taxa and fMRI features suggest potential gut-brain interactions influencing CI. For example, *Faecalibacterium* abundance was positively associated with connectivity in the salience network, while *Veillonella* showed links to executive function networks. These findings support the synergistic value of integrating multi-omics data for uncovering mechanisms underlying CI in PD.

This study demonstrates the utility of combining neuroimaging and microbiome data to enhance predictive performance and biological insight. The identified biomarkers may serve as a foundation for developing microbiome-targeted interventions and neuroimaging-guided strategies for managing cognitive decline in PD. Future work should focus on larger, longitudinal datasets and explainable AI approaches to further refine this integrative methodology.

## 1 INTRODUCTION

Parkinson’s disease (PD) is a complex neurodegenerative disorder affecting millions of people worldwide, characterized by the accumulation of alpha-synuclein (α-Syn) aggregates in neurons and the progressive development of motor and non-motor impairments ^1^. Among non-motor symptoms, cognitive impairment (CI) is particularly devastating, often progressing from mild cognitive impairment (MCI) to Parkinson’s disease dementia (PDD). The presence of CI significantly diminishes quality of life and creates significant challenges in the management of the disease, thereby imposing an absolute need for reliable biomarkers that can help with early diagnosis and therapeutic intervention ^2,3^.

Recent advances in neuroimaging, particularly resting-state functional magnetic resonance imaging (rs-fMRI), have provided insights into brain network disruptions associated with PD-related CI ^4,5^. Resting-state functional magnetic resonance imaging (rs-fMRI) studies have revealed that Parkinson’s disease (PD) is linked to profound alterations in large-scale brain networks, including the default mode network (DMN), salience network (SN), and executive function networks ^6,7^. Dysfunction in these networks, especially the weakened connectivity in the DMN and the disrupted interaction between the SN and executive function networks, has been implicated in cognitive impairment in PD, thereby pointing to the critical role played by network dysfunction in the disease pathogenesis ^8,9^.

In parallel, the gut microbiome has been considered a crucial player in the pathophysiology of neurodegenerative diseases through the gut-brain axis. Dysbiosis has been linked to neuroinflammation and altered neurotransmitter signaling, suggesting a potential role in neurodegeneration ^10^. While prior studies have independently explored neuroimaging and microbiome data, integrating these modalities remains an emerging field and could offer a more comprehensive understanding of CI mechanisms ^11,12^.

Integrating high-dimensional and multimodal data is rather challenging in clinical studies due to the limited sample size^13^. It is obvious that both rs-fMRI connectivity matrices and microbiome profiles contain a large number of features, most of which are noisy and redundant ^14,15^. Addressing these complexities necessitates the use of advanced machine learning (ML) techniques, which facilitate feature selection, dimensionality reduction, and predictive modeling ^16^. These methodologies not only enhance the interpretability of a model but also help in the identification of biologically meaningful patterns by reducing noise and eliminating redundant variables.

This study aims to integrate rs-fMRI and microbiome data to identify multimodal biomarkers predictive of CI in PD. Using a systematic feature selection pipeline, we applied linear and nonlinear ML models to evaluate the predictive value of independent and joint datasets. Our findings highlight the superior performance of integrative models, demonstrating the potential of combining neuroimaging and microbiome data to uncover key biomarkers and gut-brain interactions associated with CI in PD.

## 2 MATERIALS AND METHODS

### 2.1 Study Design and Participants

This study was approved by the Istanbul Medipol University Ethics Committee (authorization number 10840098-604.01.01-E.3958), and informed consent was obtained from all participants. Detailed information regarding the larger cohort, exclusion criteria, and rs-fMRI data can be found in our previous publication ^17^.

From the larger cohort, a subset of 60 adults aged 50–75 years with both oral and stool samples available for microbiome analysis were selected and assigned to three groups: healthy controls (HC, n = 21), mild cognitive impairment (MCI, n = 20), and Parkinson’s disease dementia (PDD, n = 19). All participants were recruited from the Neurology Clinic at Medipol Education and Research Hospital and Bakırköy Neurological and Psychiatric Diseases Research and Training Hospital. Clinical data, including cognitive assessments, were collected for all participants ^17^.

Individuals with head, trauma, stroke, exposure to toxic substances, substance use, history of antibiotic or probiotic use in the last month, chronic serious diseases (diabetes, cancer, renal failure, etc.), autoimmune diseases, smokers, pregnancy, and symptoms suggestive of Parkinson’s syndromes were excluded from the study. Demographic data, including age, gender, and years of education, were recorded and neuropsychological tests were performed during clinic visits.

### 2.2 The Resting State Functional MRI (rs-fMRI) Data Preprocessing and Feature Vectorization

rs-fMRI data preprocessing and downstream analysis was described in our previous publication ^17^. This matrix quantifies the unique functional connectivity between brain regions while controlling for the influence of all other regions, with values ranging from -1 to +1 positive values indicating a direct relationship, negative values indicating an inverse relationship, and values near zero suggesting minimal direct connectivity. To facilitate analysis, the connectivity matrix was vectorized by removing the upper or lower triangular portion (excluding the diagonal), which yielded 4950 unique connectivity features per participant. Consequently, 60x4950 dimensional fMRI feature matrix was derived for three disjoint groups and constituted the basis of subsequent group comparisons.

### 2.3 Computational Strategies for Neuroimaging and Metagenomic Data Integration and Feature Selection

#### 2.3.1 Feature Engineering and Feature Selection Applications in fMRI Dataset

We employed a stepwise feature selection process to refine the 4950 connectivity features in the training dataset. Initially, min-max normalization was applied to standardize all features. Subsequently, an Analysis of Variance (ANOVA) test was conducted to identify features with significant differences (p < 0.05) across the HC, MCI, and PDD groups, effectively eliminating irrelevant and redundant features. To control the false positive rate, False Discovery Rate (FDR) correction was applied, and features with significant p-values (p < 0.05) post-FDR correction were retained for further analysis. Then, a subset of the most discriminative features was generated from the high-dimensional feature vector using the random forest (RF) method with 10-fold cross-validation. In each iteration, the RF algorithm ranked the features according to their relative values, and the top 15 features were selected according to feature importance. The selection counts of each feature were summed and ranked, and the top 15 features with the highest selection count were selected.

To address potential multicollinearity among the remaining features, we used partial correlation coefficients. If two features had a correlation coefficient of absolute value ≥0.7 and p-value <0.05, the feature with lower importance score was removed. After all these feature selection processes, 11 features stood out from 4950 features using machine learning algorithms and further additional analyses were continued using these feature subsets.

#### 2.3.2 Feature Selection in Metagenomic Datasets (LassoCV Approach)

Feature selection for fecal and saliva microbiome datasets was performed using LassoCV regression with 10-fold StratifiedKFold cross-validation to optimize the alpha regularization parameter (Fecal ASV dataset: 0.08087; Saliva ASV dataset: 0.07034). The maximum number of iterations was set to 100,000 to ensure model convergence. Prior to training, all features were standardized, and only those with nonzero regression coefficients were retained. The initial LassoCV model identified 25 ASVs, of which 10 with positive regression coefficients were selected. Further refinement was applied by retaining features with absolute regression coefficients (|β|) greater than 0.05, resulting in a final set of 17 ASVs.

This stepwise filtering ensured that only the most informative and biologically relevant features were carried forward for downstream analyses.

#### 2.3.3 Integrating Neuroimaging and Metagenomic Data

Fecal and saliva ASVs were first integrated into a common matrix without prior feature selection following the early integration approach (Figure 1). The dataset was split into 70% training and 30% test sets. A LassoCV model with 10-fold cross-validation was applied, optimized using an iterative solver with a maximum of 100,000 iterations for convergence, and features with nonzero coefficients were retained after optimization (alpha = 0.08254). Then, the metagenomic features obtained here and 11 features obtained from the fMRI dataset, where feature selection was applied independently, were combined with the middle integration. The matrix we used for the joint model contains a total of 44 features (fMRI: 11, Fecal ASVs: 18, Saliva ASVs: 15).

**Figure 1.**
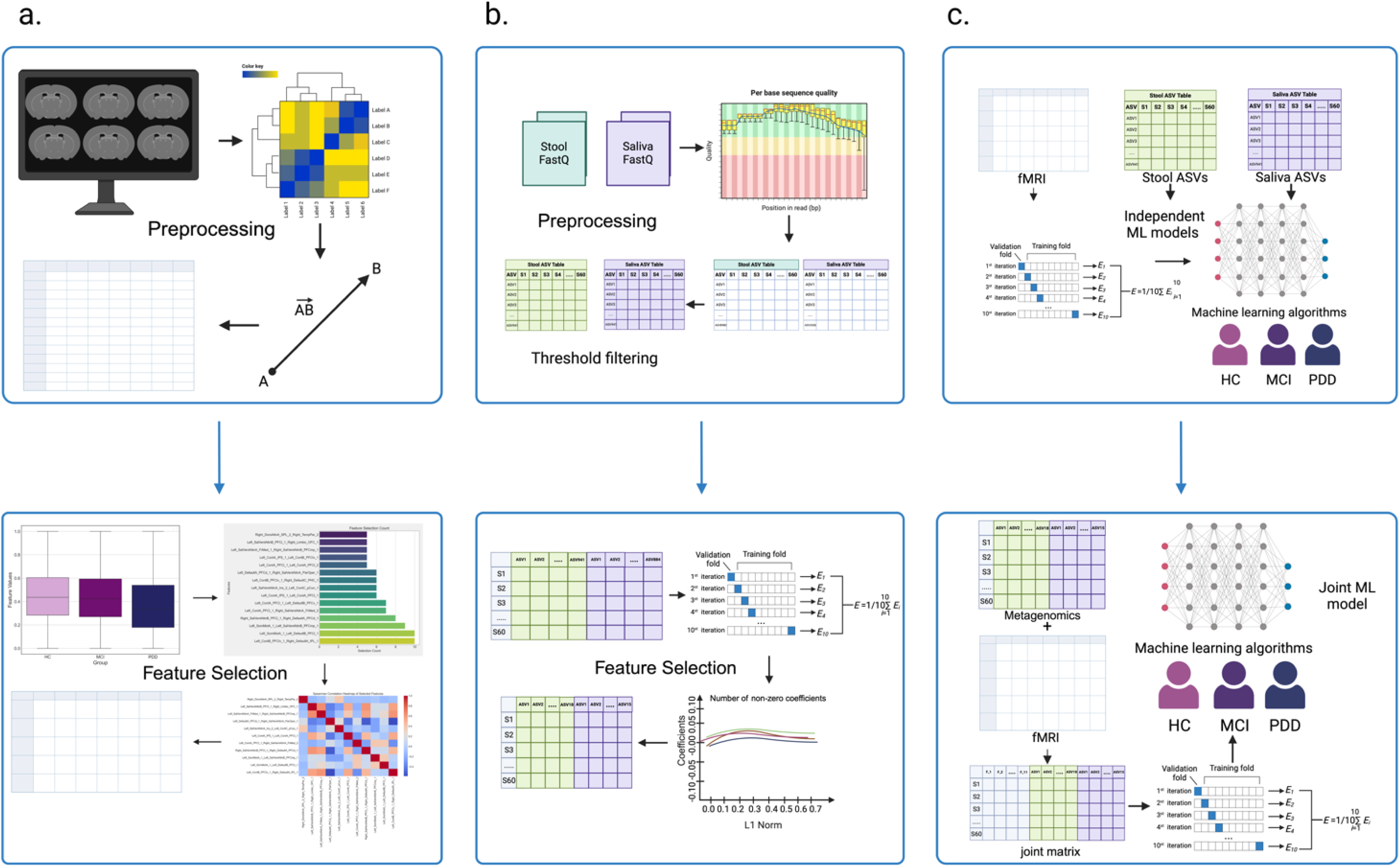
Flowchart of the machine learning process ^71^. a. The rs-fMRI scans were preprocessed, and then a total of 4950 features were extracted. There methods, including ANOVA (Analysis of Variance) test, random forest, and removing features with high autocorrelation, were used to select the characteristics with high predictive performance. b. Preprocessing and filtering steps were applied to the sequence files (FastQ) of the samples taken from the Stool and Saliva microbial regions. The resulting Stool ASV and Saliva ASV tables were used for independent ML models. Metagenomic datasets (Stool ASVs and Saliva ASVs) were integrated into a joint matrix without applying any feature selection method, following an early integration approach. StandardScaler was used to standardize features, and a LassoCV model was used for feature selection and regularization. c. The obtained fMRI, Stool ASV and Saliva ASV Datasets were also applied to the algorithms as independent ML models. The metagenomic and fMRI features were combined with middle integration. Finally, multiple machine learning models (Random forest, SVM, DT, AdaBoost, XGBoost, Logistic regression) were built to predict patients with HC, MCI and, PDD.

### 2.4 Machine Learning on Neuroimaging and Metagenomic Datasets

#### 2.4.1 Independent Models and Joint Model

The dataset was partitioned into training (70%) and validation (30%) sets using train_test_split. To guarantee methodological rigor and prevent data leaking, the same training split obtained from this technique was utilized for both feature selection and ML model training.

To assess potential confounding effects, we examined differences in metadata across study groups. Statistical analyses revealed significant differences in age and education level, while no significant variation was observed for sex. To account for these potential confounders, models were additionally trained with age and education level as features. However, the inclusion of these variables did not result in statistically significant changes in model performance metrics, indicating that their confounding effects were negligible.

For independent and joint models, machine learning classifiers were applied, including random forest, decision tree, support vector machine (SVM), AdaBoost, XGBoost, and logistic regression. These models were selected to accommodate both linear and nonlinear data structures, ensuring a robust comparison of predictive performance.

#### 2.4.2 Hyperparameter Optimization and Performance Evaluation

Hyperparameter optimization was performed using GridSearchCV within the training set, ensuring systematic exploration of predefined parameter spaces. The best hyperparameter configurations were identified and subsequently used to train the final models.

Model performance was assessed using 10-fold cross-validation, ensuring balanced class distribution across folds. The evaluation metrics included accuracy, precision, recall, F1-score, and ROC-AUC score, providing a robust assessment of model effectiveness.

### 2.5 Bioinformatics and Statistical Analyses

Bioinformatics and statistical analyses were conducted using R (v3.6.1), Python (v3.11.1), and Java (v11.0.17). Machine learning analyses were implemented using scikit-learn (v1.5.1) and related libraries. Feature selection and model training were performed using StratifiedKFold cross-validation and GridSearchCV for hyperparameter tuning.

For statistical comparisons, one-way analysis of variance (ANOVA) was used for parametric data, while the Kruskal-Wallis test was applied for non-parametric data. When significant differences were detected, the Mann-Whitney U test was applied as a post-hoc test to determine comparisons between groups.

All p-values were adjusted for multiple testing using the Benjamini-Hochberg method, where appropriate. Multivariate statistical methods were calculated according to the selected models. We applied the SparCC algorithm ^18^with 20 iterations to extract linear relationships between the features selected by the ML algorithms in our fMRI, Stool ASVs and Saliva ASVs datasets. Additionally, we included a bootstrap process with 99 resampling iterations the boot_cor() function to estimate correlations. The obtained results were then plotted using the Gephi platform (version: 0.10.1; Java: 11.0.17). In addition, the Louvain algorithm ^19^was added to detect communities in the networks. Information on the topological properties of each network we created is also available in the supplementary materials. The pheatmap, lattice and ggplot2 R packages were also used for visualizations.

## 3 RESULTS

Integration of rs-fMRI and microbiome data revealed significant associations between disruptions in brain connectivity and specific microbial features linked to cognitive impairment severity in Parkinson’s disease. Notably, machine learning analyses accurately classified participants across healthy controls (HC), mild cognitive impairment (MCI), and PD dementia (PDD), highlighting predictive biomarkers in both microbiome composition and functional brain networks (Figure 1). Before detailing these findings, we summarize the demographic and clinical characteristics of participants in Table 1.

**Table 1.** Comparison of participants metadata features between groups. Statistical significance was checked by the Kruskal-Wallis test followed by Mann-Whitney U tests as post-hoc analysis.

| Metadata | HC<br>(mean ±<br>std) | MCI<br>(mean ±<br>std) | PDD<br>(mean ±<br>std) | Kruskal-Wallis<br>H-value | p-value | FDR-corrected<br>p-value | Mann-Whitney<br>U (HC vs<br>MCI) | p-value<br>(HC<br>vs<br>MCI) | Mann-Whitney<br>U<br>(HC vs<br>PDD) | p-value<br>(HC<br>vs<br>PDD) | Mann-Whitney<br>U (MCI<br>vs PDD) | p-value<br>(MCI vs<br>PDD) |
| --- | --- | --- | --- | --- | --- | --- | --- | --- | --- | --- | --- | --- |
| Age | 58.4762<br>± 8.4239 | 66.4500<br>± 8.9764 | 71.8421<br>± 8.1258 | 18.9926 | 0.0001 | 0.0001*** | 99.0000 | 0.0039 | 51.0000 | 0.0001 | 121.5000 | 0.0554 |
| Gender | 0.6667 ±<br>0.4830 | 0.3000 ±<br>0.4702 | 0.4737 ±<br>0.5130 | 5.4333 | 0.0661 | 0.0661 | 287.0000 | 0.0212 | 238.0000 | 0.2295 | 157.0000 | 0.2787 |
| Education | 10.5714<br>± 5.1046 | 6.6500 ±<br>4.0167 | 4.4737 ±<br>4.5628 | 15.6266 | 0.0004 | 0.0005*** | 300.0000 | 0.0146 | 329.5000 | 0.0003 | 256.5000 | 0.0475 |
| MMSE | 27.9048<br>± 1.7001 | 23.6000<br>± 3.6907 | 19.1579<br>± 3.2363 | 41.3516 | 0.0000 | 0.0000*** | 382.5000 | 0.0000 | 399.0000 | 0.0000 | 324.0000 | 0.0002 |
| CDRS | 0.0000 ±<br>0.0000 | 0.5000 ±<br>0.0000 | 1.1053 ±<br>0.3153 | 58.4101 | 0.0000 | 0.0000*** | 0.0000 | 0.0000 | 0.0000 | 0.0000 | 0.0000 | 0.0000 |

Statistically significant differences between groups are shown in the relevant columns (HC vs MCI, HC vs PDD, MCI vs PDD). The significance levels were defined as follows: \* $p < 0.05$ , \*\* $p < 0.01$ , and \*\*\* $p < 0.001$ .

### 3.1 Demographic and Clinical Characteristics of Participants

Demographic profiles differed significantly across HC, MCI, and PDD groups (p < 0.0001). A progressive decline in cognitive function was observed from HC to PDD, reflected by decreasing MMSE scores and increasing CDRS scores (both p < 0.0001). Additionally, the PDD subjects were older and less educated than the HC and MCI groups (p < 0.001). Importantly, there was no significant difference in sex distribution among groups (p = 0.0661), ensuring balanced gender representation across study cohorts. Pairwise comparisons also determined significant cognitive differences to support a definite trend of cognitive decrease from HC to MCI to PDD (Table 1).

### 3.2 Performance Evaluation of Machine Learning Methods on rs-fMRI Dataset

In this study, various machine learning models were evaluated for their performance on rs-fMRI data (Table S2.1). On the training set (n = 42), XGBoost achieved perfect metrics (accuracy, F1 score, AUC = 1.0) but showed overfitting, with validation accuracy dropping to 0.556. Similarly, SVM demonstrated high training accuracy (0.952) but performed poorly on the validation set (accuracy = 0.5). Random Forest (RF) displayed better generalizability, maintaining a validation accuracy of 0.833, while Logistic Regression offered stable but moderate performance (accuracy = 0.556).

In terms of discriminatory power (Figure S2.1), RF achieved the highest average AUC (0.94), excelling in differentiating the MCI group (AUC = 1.0). XGBoost and Logistic Regression had average AUC scores of 0.81, reflecting good class discrimination, while SVM and AdaBoost showed moderate AUCs (0.76 and 0.78). The Decision Tree model exhibited the lowest AUC (0.59), indicating weaker classification performance. These findings suggest that while complex models like RF and XGBoost excel in capturing detailed patterns, simpler models like Logistic Regression provide more consistent generalization.

### 3.3 Machine Learning Model Performance on the Microbiome Datasets

#### 3.3.1 Stool ASVs Dataset

The performance of various machine learning models was evaluated using the stool ASVs dataset (Table S2.2). Logistic Regression performed exceptionally well on the training dataset, n = 42, with perfect accuracy, F1 score, and AUC = 1.0 metrics. It generalized very well to the validation dataset, n = 18, with accuracy 0.833. Random Forest performed well on the training set with an accuracy of 0.786 and AUC of 0.937; however, signs of overfitting, as its validation accuracy dropped to 0.611. On the other hand, the SVM displayed moderate training performance (accuracy = 0.524) but performed better on the validation set (accuracy = 0.667, AUC = 0.75), suggesting stronger generalization than Random Forest.

AdaBoost and XGBoost demonstrated moderate training accuracies (0.595 and 0.643, respectively), but their validation performance declined significantly (0.556 and 0.444, respectively). The Decision Tree model had a training accuracy of 0.667 but a poor validation accuracy of 0.5, making it the least effective approach. (Figure S1.2). ROC curve analysis identified Random Forest as the best model, with the highest average AUC (0.79), while its performance for the HC class was remarkably higher, with an AUC of 0.90.

Logistic Regression also performed well, achieving an AUC of 0.78 overall and perfectly classifying the HC group (AUC = 1.0). For the SVM, this yielded average AUCs: 0.75; highest for the class PDD: AUC = 0.92 (Figure S2.2). In contrast, the Decision Tree and XGBoost had the worst averaged AUCs: both scored 0.64 for the poor class discrimination value. Overall, the best generalizations had been achieved with Random Forest and Logistic Regression: RF was really robust, while Logistic Regression turned out exceptionally good for the classification of HC.

#### 3.3.2 Saliva ASVs Dataset

For the saliva dataset (Table S2.3), Random Forest, SVM, and XGBoost achieved perfect training accuracy, indicating strong model fit. However, all three models exhibited overfitting, as their validation performance dropped significantly. RF exhibited a significant drop in validation performance, highlighting overfitting, while SVM had low generalizability (validation accuracy = 0.444, AUC = 0.639).

In contrast, Logistic Regression and AdaBoost provided more balanced across training and validation. Although their predictive power was moderate, their results were more stable. The Decision Tree model is the least effective among these models because of its simpler structure.

ROC curve analysis showed that Random Forest was best for the MCI class (AUC = 0.96), and on average, has the highest AUC (0.74). SVM and XGBoost performed moderately well with XGBoost achieving the highest classification performance for PDD (AUC = 0.79). Logistic Regression and AdaBoost have more stable average AUCs of 0.75 and 0.72, respectively (Figure S2.3).

The ensemble methods, such as RF and XGBoost, performed well in training but they suffered from overfitting; on the other hand, simpler models like Logistic Regression provided more stable results, making them potentially more reliable for the Saliva ASV dataset.

### 3.4 Assessing the Performance of Machine Learning Models for a Joint Model

The joint model, integrating features from rs-fMRI with stool and saliva ASVs, demonstrated an excellent performance across multiple machine learning approaches. Within the validation data set (n = 18), RF performed best with an accuracy of 0.889 and AUC of 0.972, reflecting outstanding generalization capability (Table S2.4).

XGBoost also performed well with an accuracy of 0.722 and AUC of 0.921. SVM and logistic regression presented balanced results, with accuracy of 0.778 and AUC of 0.866 and 0.852, respectively. Decision Tree and AdaBoost had the lowest classification performance, with Decision Tree yielding the lowest validation accuracy (0.667) and AUC (0.84).

ROC curve analysis revealed RF to have the best discriminatory ability, with the highest mean AUC (0.97), perfectly classifying MCI (AUC = 1.00) and nearly perfect classification of PDD (AUC = 0.96). XGBoost performed well, with a mean AUC value of 0.92, while SVM and Logistic Regression provided stable yet somewhat lower discrimination (AUC = 0.87 and AUC = 0.85, respectively) (Figure S2.4). AdaBoost and Decision Tree had moderate performance, with poor efficiency for Decision Tree, as it is based on a simpler structure. Overall, Random Forest and XGBoost were the most effective models for integrating multimodal datasets, demonstrating their ability to capture complex patterns in cognitive impairment. Simpler models, such as Logistic Regression and Decision Tree, provided steady but less detailed predictions.

### 3.5 Comparison of Joint and Independent Models

#### 3.5.1 Model Performance Comparison

The integrated model outperformed the independent models, achieving higher accuracy and AUC scores across most machine learning algorithms (Figure 2.1.a., Figure 2.2.a.). In the joint model, the best performances were from Random Forest (accuracy = 0.889, AUC = 0.972) and Logistic Regression (accuracy = 0.778, AUC = 0.852).

**Figure 2.1.**
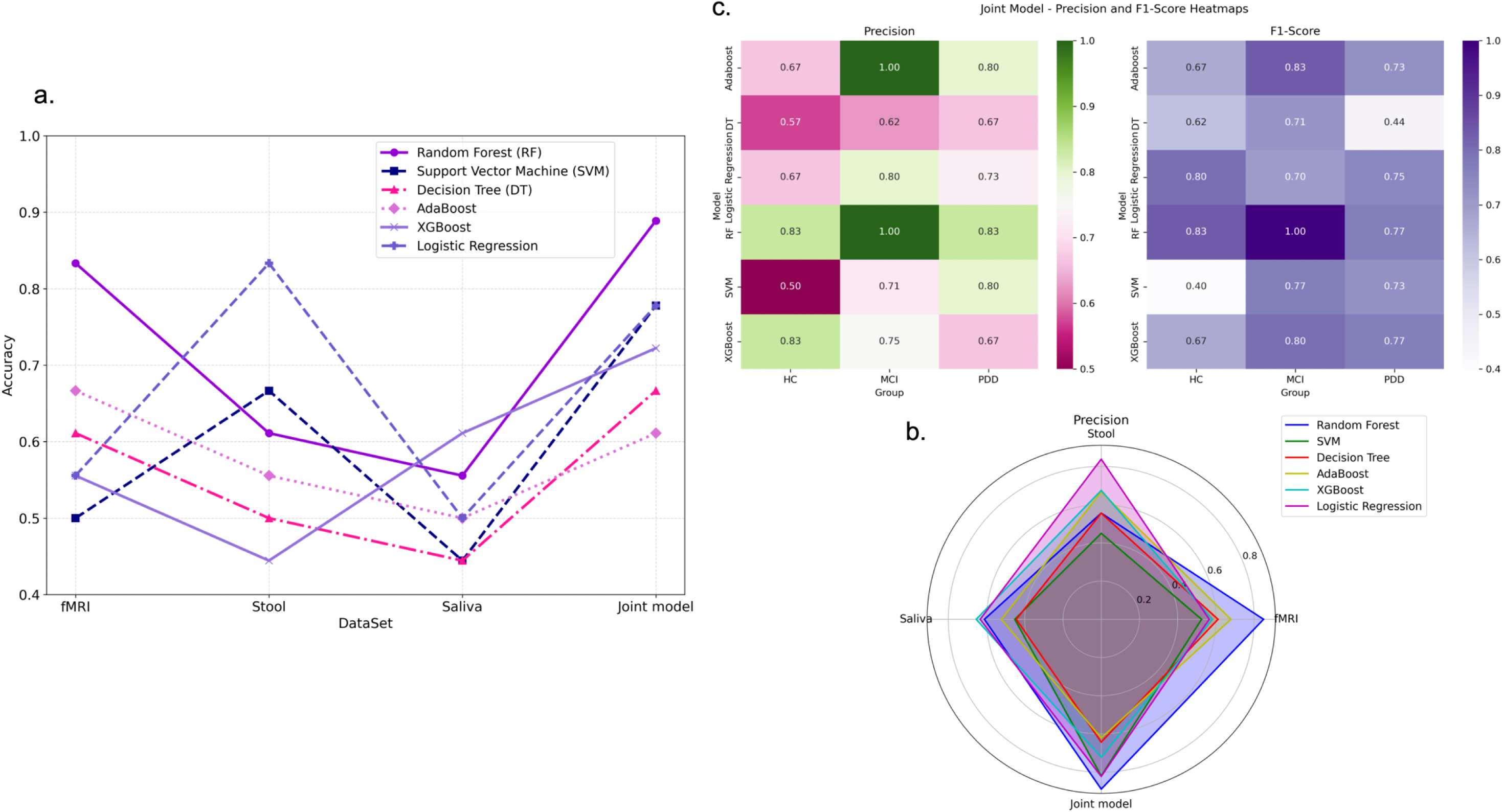
Performance comparison of machine learning algorithms between the joint model and independent models. a. The line plot illustrates the accuracy performance comparisons between the joint model and the independent models across various machine learning algorithms. b. The radar chart shows the precision comparisons between the joint model and the independent models among various machine learning algorithms. c. The precision and F1-score comparisons of the joint model across study groups using different machine learning algorithms are presented as heatmaps.

**Figure 2.2.**
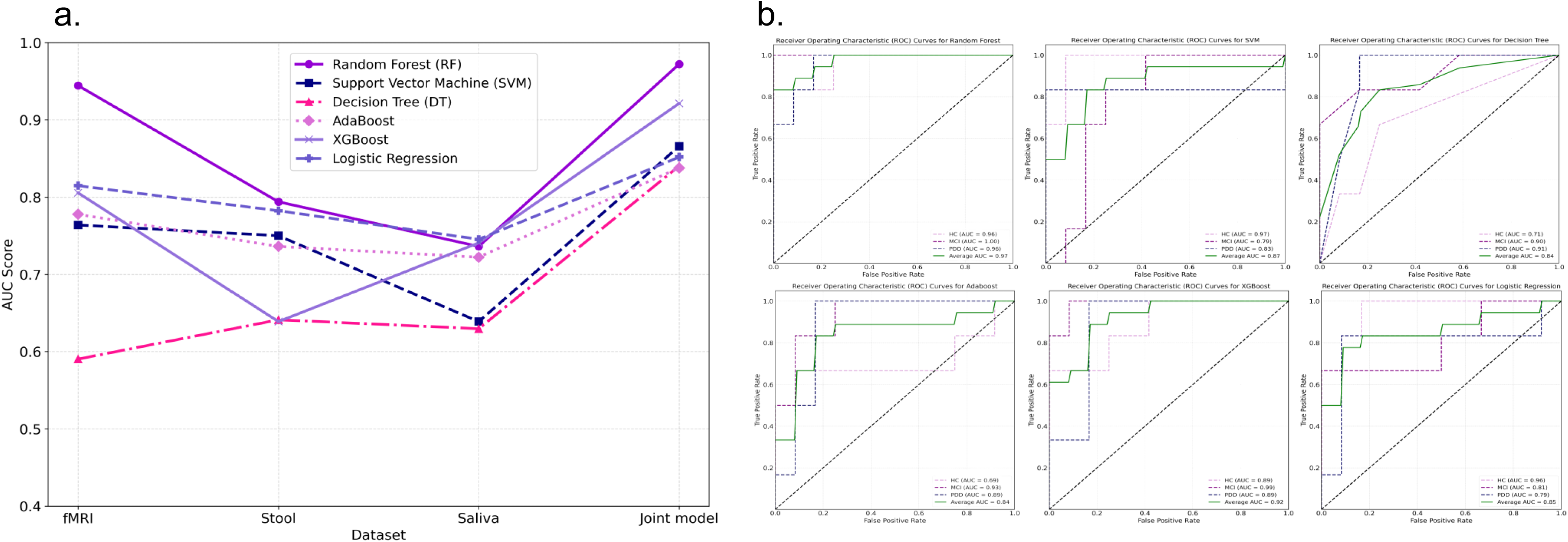
Performance comparison of machine learning algorithms between the joint model and independent models. Comparative analysis of the performance of the joint model across study groups. a. The line plot illustrates the AUC score performance comparisons between the joint model and the independent models across various machine learning algorithms. b. Illustrates the area under the ROC curve (AUC) for the joint model across different machine learning algorithms. AUC scores are provided for the study groups (HC, MCI, PDD) included in the model, demonstrating its performance in distinguishing between these groups.

**Figure 2.3.**
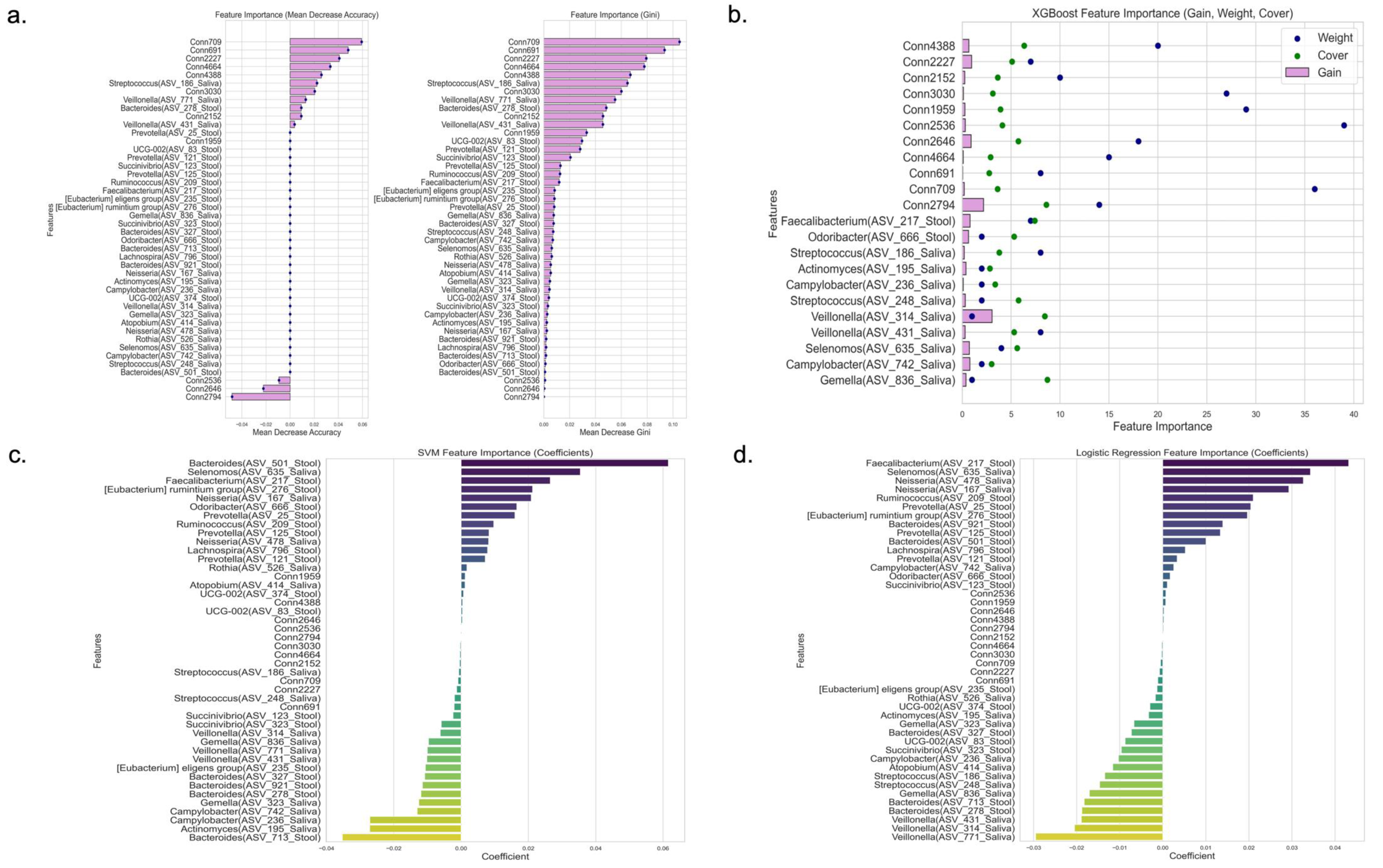
Comparative analysis of feature importance across four different machine-learning algorithms that showed better performance in our study. Panels (a) through (d) show feature importance ranking for selected features using four different algorithms: Random Forest (a), XGBoost (b), Support Vector Machine (SVM) (c), and Logistic Regression (d).

In contrast, performance among individual models was diverse. Logistic Regression gave good results with the Stool dataset (accuracy = 0.833), but in the case of Saliva, the result was poor (accuracy = 0.5) (Figure 2.1.a., Figure 2.2.a.). Similarly, Support Vector Machines (SVM) and AdaBoost showed better generalization capabilities on the Stool dataset compared to the Saliva dataset.

The integrated model performed better generally with respect to precision and F1 scores, with Random Forest achieving the highest (precision = 0.889, F1 score = 0.889), indicating a well-balanced and robust classification ability (Figure 2.1.b.,c.) Independent models showed unstable performances, but they appeared to be drastically improved in precision and F1 scores when applied to the joint set as a whole, especially for XGBoost and Logistic Regression.

The integrated approach reduced performance variability, enhanced robustness, and provided a more reliable classification framework, emphasizing the importance of multimodal data integration for cognitive impairment assessment in Parkinson’s disease.

#### 3.5.2 Feature Importance in the Joint Model

Feature importance analysis in the joint model revealed that both neuroimaging (fMRI) and microbiome datasets (stool and saliva ASVs) contributed significantly to classification accuracy across machine learning models, including Random Forest, XGBoost, Logistic Regression, and SVM (Figure 2.3).

The fMRI dataset showed that the areas related to core brain networks, specifically the dorsal attention network, salience-attention network, default mode network, and control network, maintained a constantly high importance across the models. The most informative features, such as Conn4388 (Dorsal Attention Network to Temporoparietal connection) and Conn2227 (Salience Network to Limbic System connection), showed significant predictive power. These areas were ranked high in Random Forest (Average Decay Accuracy = 0.0259, 0.0407, respectively) and XGBoost tests (Gain = 0.687 and 0.973, respectively), as well as yielding large coefficients in Logistic Regression and Support Vector Machine (SVM) tests.

Additional critical features included connectivity within the salience network, Conn2152 (Salience Network connection), have also highlighted the role of attention-related mechanisms in differentiating degrees of cognitive impairment. It was especially clear in XGBoost (Gain = 0.273) and Random Forest (Average Decrease Accuracy = 0.0093), which outlines a potent predictor.

Analogously, the default mode-control network interconnectivity, Conn3030 (Default Mode to Salience Network connection) and Conn2536 (Control Network connection), demonstrated their cooperative role in the classification task, indicating some relation of those networks to neurodegenerative changes. The Random Forest model further highlighted regions related to the somatomotor network, such as Conn709 (Somatomotor to Default Mode Network connection), and areas connected with salience and attention, bringing out their importance in the discrimination between healthy and pathological states. These findings underline that neural connectivity alterations within these networks are of paramount importance for determining cognitive impairment in Parkinson’s disease.

In the stool dataset, the microbial taxa *Faecalibacterium* (ASV_217), *Odoribacter* (ASV_666) and *Bacteroides* (ASV_278) were consistently identified as the most important predictors. *Faecalibacterium*, with its known anti-inflammatory characteristics, was of considerable importance in both Random Forest (Average Decay Accuracy = 0.0432) and XGBoost (Gain = 0.815), thus establishing its potential value as a biomarker of neurodegenerative disease progression. *Odoribacter and Bacteroides* also showed strong predictive contributions, particularly in Random Forest and Logistic Regression models, thus supporting their use as markers of various health states.

In the saliva dataset, some of the notable taxa included *Streptococcus* (ASV_186), *Veillonella* (ASV_771), and *Selenomonas* (ASV_635), which were continuously identified across different models. For instance, *Streptococcus* showed high importance in Random Forest (Gini Significance = 0.065) and XGBoost (Gain = 0.211). Besides, *Veillonella* and *Selenomonas* exhibited high predictive powers, particularly in Logistic Regression, which brought forth the unique role oral microbiota could play in identifying disease-specific indicators.

Collectively, results provide that neural network alterations, particularly in attention and salience networks, play a key role in cognitive impairment classification. The complementary role of microbiome-derived features, especially in stool and saliva samples, further strengthens the importance of integrating neuroimaging and microbiome data to improve classification accuracy and enhance our understanding of Parkinson’s disease progression.

### 3.6 Comparison of Features Selected by Machine Learning Algorithms Across Study Groups

#### 3.6.1 rs-fMRI Dataset

The Kruskal-Wallis and Mann-Whitney U tests were applied to assess significant differences in functional connectivity across the HC, MCI, and PDD groups, representing different stages of cognitive impairment (Figure 3). Several key rs-fMRI features showed statistically significant variations between groups, highlighting their potential as biomarkers. Conn4388 (Dorsal Attention A to Temporoparietal Network connection) and Conn2536 (Control Network connection) showed high Kruskal-Wallis H values (111.31 and 187.815, respectively) with p-values <0.005, indicating significant differences in across groups.

**Figure 3.**
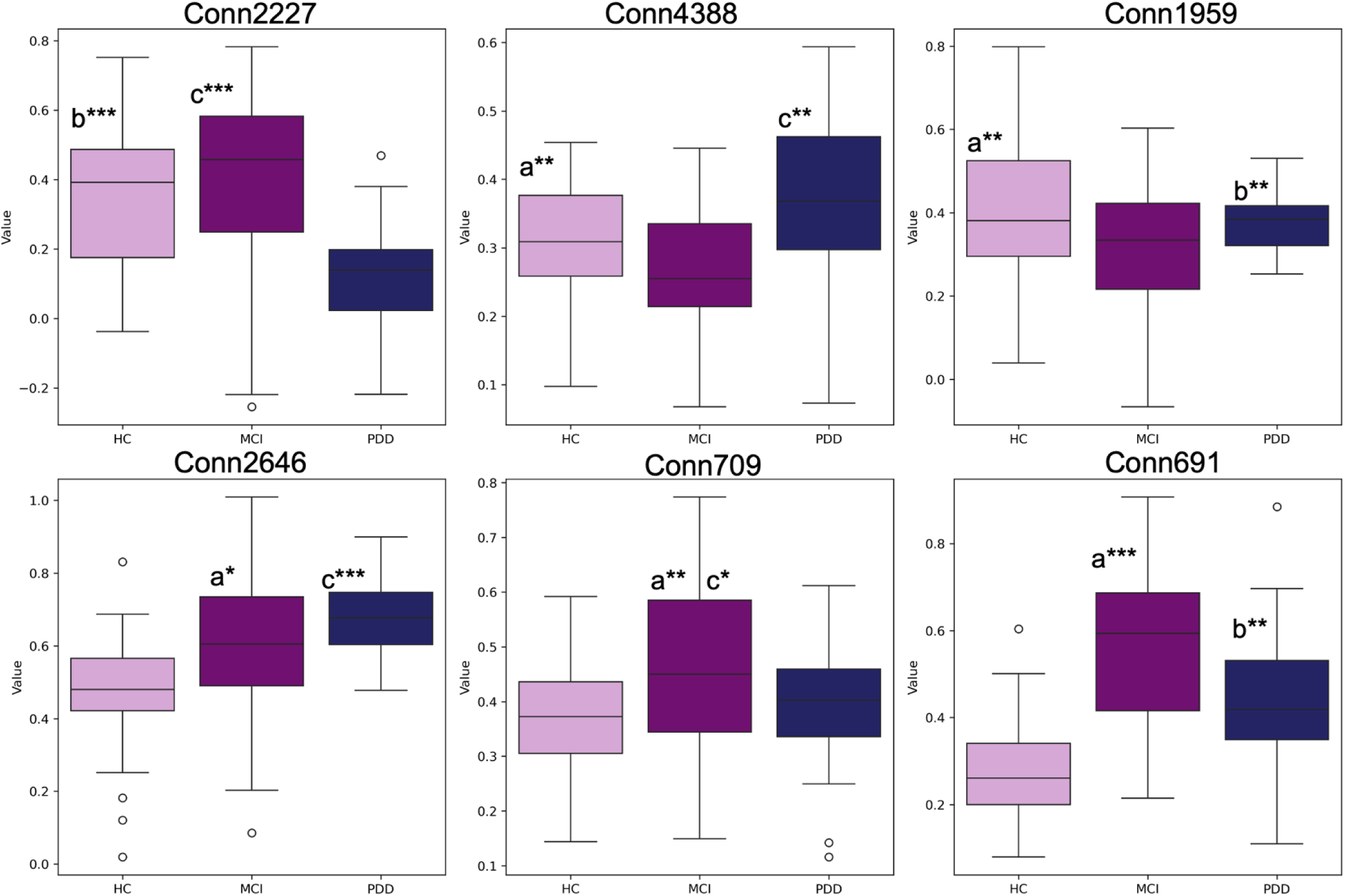
Inter-group comparison of 6 prominent features selected by ML algorithms from the rs-fMRI dataset. Statistical significance was checked by the Kruskal-Wallis test followed by Mann-Whitney U tests as post-hoc analysis. Letters indicate statistically significant differences between groups: ’a’ for significant differences between HC and MCI, ’b’ between HC and PDD, and ’c’ between MCI and PDD. The number of asterisks represents the level of significance (*p < 0.05, **p < 0.01, ***p < 0.001).

Conn2227 (a Salience Network to Limbic System connection) showed significant group-wise differences, particularly between HC and PDD (U = 3.230, p = 0.0009) and between MCI and PDD (U = 3.230, p = 0.0002), suggesting its relevance in distinguishing more advanced stages of disease progression. Similarly, Conn2646 (a Control Network to Salience Network connection) showed significant differences between HC-MCI (U = 1.120, p = 0.011) and HC- PDD (U = 570, p = 0.0001), further supporting its role in tracking early to late-stage cognitive changes.

Among the identified features, Conn2794 (a Control to Default Mode Network connection), demonstrated the highest Kruskal-Wallis value of H (222.097, p < 0.0001) and showed strong differences between HC and PDD (U = 3.510, p < 0.0001) and MCI and PDD (U = 3.350, p < 0.0001). The strong ability of this feature to differentiate between healthy, early, and advanced disease states underscores its potential as a critical biomarker for neurodegenerative progression.

#### 3.6.2 Metagenomic Datasets

Microbial composition varied significantly across HC, MCI, and PDD groups, with distinct shifts in both gut and oral microbiota that progressed with neurodegeneration (Figure 4). In the fecal microbiota, the relative abundance of *Bacteroides* species (e.g., ASV_278, ASV_327) were highly elevated in PDD, while beneficial taxa such as *Faecalibacterium* (ASV_217) and *Lachnospira* (ASV_796) significantly decreased in abundance with disease progression. Similarly, *Odoribacter* and *Prevotella* decreased in their relative abundance, with the most pronounced reduction in the PDD group, reflecting a shift toward the dysbiotic gut microbiota profile (Figure 4.a.).

**Figure 4.**
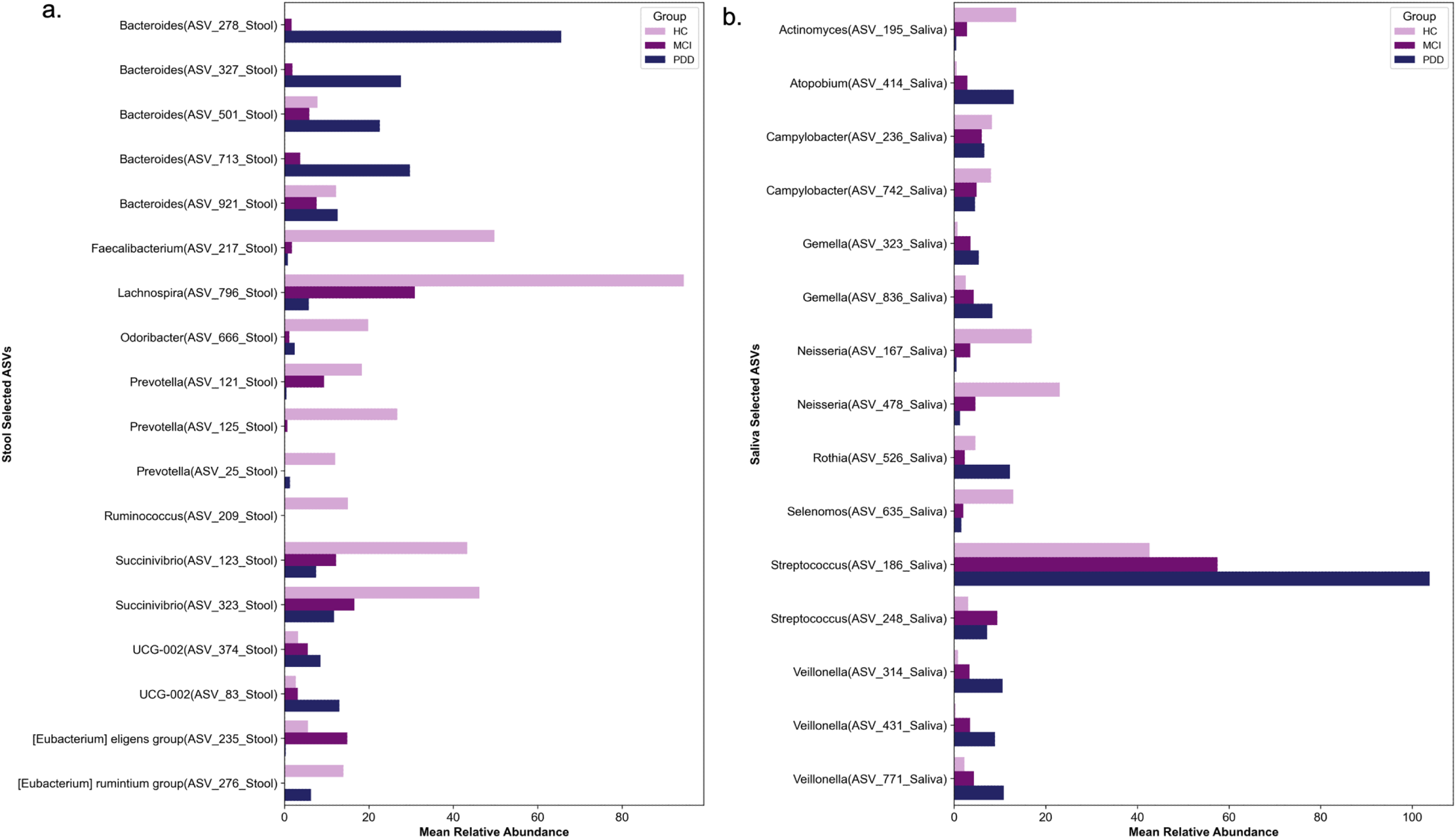
Comparative analysis of inter-group mean relative abundances for features selected by machine learning algorithms from Stool and Saliva ASV datasets. Mean relative abundance of (a) Stool ASVs and (b) Saliva ASVs in between different study groups (HC, MCI, PDD).

Salivary microbiota followed a similar trend, with protective taxa such as *Actinomyces* and *Neisseria*, decreased from HC to PDD, whereas the opportunistic taxa *Atopobium, Gemella, Streptococcus,* and *Veillonella* were increased in PDD, suggesting their potential association with disease progression (Figure 4.b.).

These findings underscore the critical role of gut and oral microbiota shifts in Parkinson’s disease progression. The observed microbial changes align with a broader pattern of neurodegenerative disease-associated dysbiosis, where a reduction in beneficial species and an increase in opportunistic microbes may contribute to or reflect disease pathology. The identification of these taxa as potential biomarkers underscores the significance of microbiome alterations in neurodegeneration.

### 3.7 Associations Between Selected ASVs and rs-fMRI Features

Spearman correlation analysis identified significant associations (|r| ≥ 0.30, p < 0.05) between microbial taxa and fMRI features, providing insights into putative gut-brain and oral-brain interactions (Figure 5.1.a.).

**Figure 5.1.**
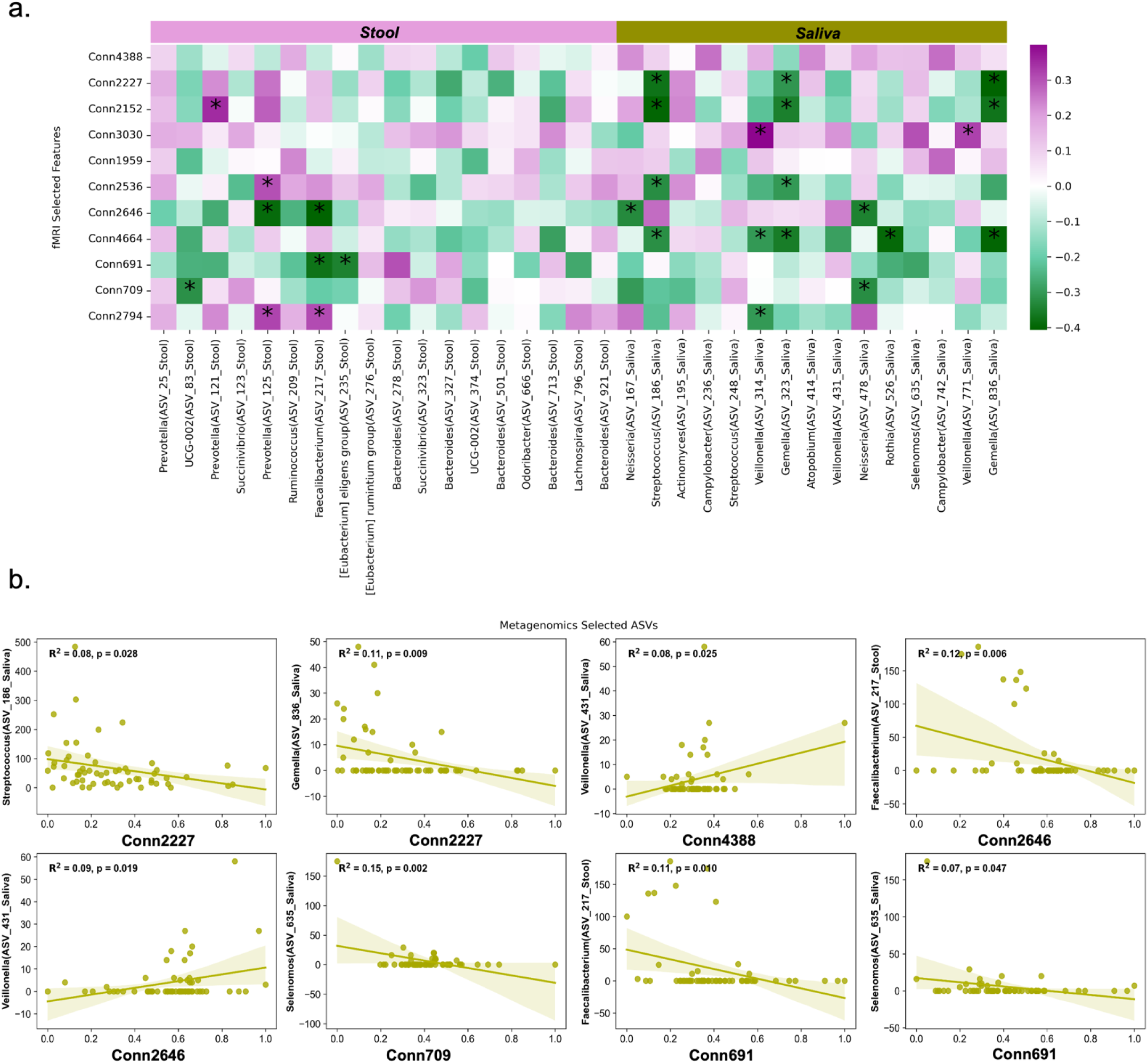
Associations between specific ASVs from fecal and saliva datasets and fMRI- selected features. a. Heatmap of Spearman’s correlation analysis between fMRI-selected features and the metagenomic ASVs. Correlation coefficients are color-coded, from dark green showing negative correlations to dark purple showing positive correlations. Statistically significant correlations between fMRI-selected features and metagenomic ASVs with |r| ≥ 0.30 and p < 0.05 are asterisked (*) to highlight meaningful associations within the dataset. b. Significant linear regressions between fMRI-selected features and specific ASVs from the stool and saliva datasets. Each scatter plot shows a linear regression line (in green) with a 95% confidence interval. R² values and p-values represent the strength and significance of each association, respectively. Associations with a p-value < 0.05 are highlighted to emphasize the statistically significant correlations.

The primary observations indicate favorable correlations between *Prevotella* (ASV_121, Stool) correlated positively with Conn2646 (a Control A to Salience Network connection; r ≈ 0.30), as well as between *Campylobacter* (ASV_236, Saliva) and Conn4388 (a Dorsal Attention A to Temporoparietal Network connection; r ≈ 0.30), implying that microbial elements may influence cognitive regulation and attentional networks. Conversely, negative correlations were detected for UCG-002 (ASV_83, Stool) with Conn2646 (a Control A to Salience Network connection; r ≈ -0.32) and for *Bacteroides* (ASV_713, Stool) with Conn709 (a Somatomotor A to Default B Network connection; r ≈ -0.34), indicating possible inverse relationship with somatomotor and control networks.

Salivary taxa also showed significant correlations. *Veillonella* (ASV_314) correlated negatively with Conn2794 (a Control B to Default A Network connection;r ≈ -0.35), and *Neisseria* (ASV_167) with Conn4664 (a Salience Network to Default A Network connection;r ≈ -0.30). These results suggest salivary taxa are associated with default mode and salience-attention network connectivity.

The scatter plots revealed significant linear relationships (Figure 5.1.b.), such as the negative associations between *Streptococcus* ASV_186 (Saliva) and Conn2227 (R² = 0.08, p = 0.028), as well as between *Faecalibacterium* ASV_217 (Stool) and Conn2646 (R² = 0.12, p = 0.006). Furthermore, positive associations were also observed for *Veillonella* ASV_431 (Saliva) with Conn2646 (R² = 0.09, p = 0.019), and *Selenomonas* ASV_635 (Saliva) with Conn709 (R² = 0.15, p = 0.002).

Network analysis using SparCC (|r| > 0.75, p < 0.05) further highlighted distinct connectivity patterns across study groups (Figure 5.2). In the HC group, well-regulated interactions were observed, such as negative correlations of *Succinivibrio* (ASV_123) with Conn2646 (a Control A to Salience Network connection; r = -0.92), suggesting microbial abundance and brain connectivity are linked.

**Figure 5.2.**
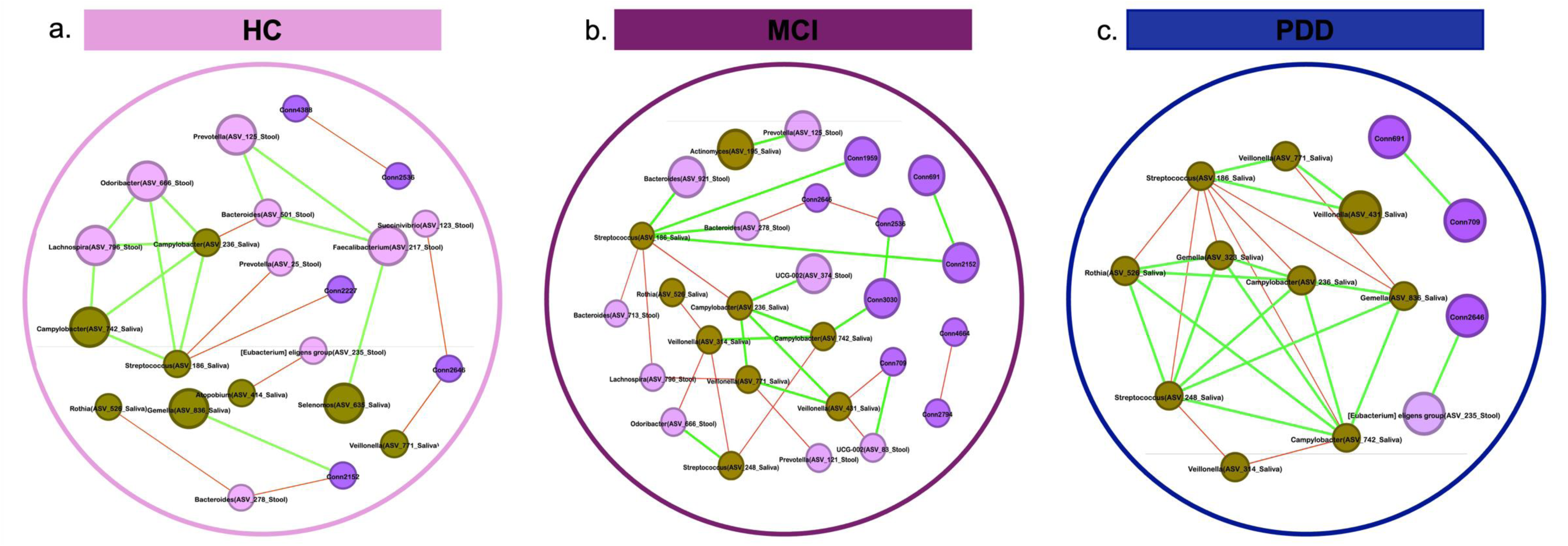
Co-abundance networks of fMRI-selected features and metagenomics-selected ASVs in different study groups. a. (HC), b. (MCI), and c. (PDD) figures study groups represent the network of respectively. Each node’s size is determined by its correlation of features. The nodes forming the network are separated according to the data sets and distinguished with different colors (fMRI: purple, Stool: lilac, Saliva: green). Only strong and significant SparCC correlations (|r| > 0.75; p<0.05) are represented as edges. Each edges size is determined by its correlation of connections, with colors indicating positive (green) and negative (red) correlations between nodes

In the MCI group, UCG-002 (ASV_83, stool) correlated positively with Conn709 (a Somatomotor A to Default B Network connection; r = 0.96), indicating microbial associations with motor and default mode regions. The PDD group exhibited fewer significant correlations, except for a strong positive association between *[Eubacterium] eligens* (ASV_235) and Conn2646 (a Control A to Salience Network connection; r = 0.84), which reflects the altered gut-brain axis interactions in advanced disease stages. These findings suggest that microbial- brain connectivity relationships shift throughout disease progression.

### 3.8 Group-Wise Comparisons of Metadata, Neuroimaging, and Microbiome Features

Demographic profiles significantly differed across HC, MCI, and PDD groups. A progressive decline in cognitive function was observed from HC to PDD, accompanied by increasing age and decreasing educational attainment (all p <0.001). Pairwise comparisons confirmed significant differences between groups, indicating a clear cognitive decline trajectory from HC to PDD. Notably, there was no significant difference between groups regarding sex distribution (p = 0.0661), supporting balanced gender representation across study groups (Table 1).

Neuroimaging features exhibited significant associations with cognitive scores. Connectivity between Control and Salience Networks was negatively correlated with MMSE scores (r = - 0.41, p = 0.01), while connectivity between Default Mode and Salience Networks was positively correlated with CDRS scores (r = 0.14, p = 0.003) (Figure S3.1).

Microbial composition was also linked to cognitive and demographic parameters. *Bacteroides* abundance was positively correlated with CDRS (r = 0.32, p = 0.04), while *Streptococcus* and *Veillonella* abundances were associated with stronger cognitive impairment. *Lachnospira* abundance was positively correlated with education level (Figure S3.1).

Regression models highlighted complex relationships between neural connectivity, microbial features, and cognitive measures. Connectivity within the Control-Awareness Network was inversely associated with MMSE scores (R² = 0.09, p = 0.03), whereas *Neisseria* abundance was positively associated with higher MMSE scores (R² = 0.12, p = 0.004) (Figure S3.2).

## 4 DISCUSSION

This study presents a novel approach to characterizing cognitive impairment in PD by integrating resting-state rs-fMRI and microbiome profiling through machine learning. While a few studies have explored the relationship between neuroimaging and microbiota ^11,12,20^, studies applying a comprehensive machine learning framework for multimodal data integration remain scarce. The systematic feature selection pipeline employed here minimizes noise, improves interpretability, and enhances predictive accuracy, demonstrating the value of integrating multi-omics data for neurodegenerative disease classification. Our multimodal framework identifies unique brain-microbiome interactions that distinguish HC, MCI, and PDD.

### 4.1 Advancing Multimodal Integration in PD Research

In this study, we integrated neuroimaging and metagenomic data to better understand the complex interplay underlying Parkinson’s disease (PD) and cognitive impairment (CI). Such integration is challenging due to the high dimensionality and inherent noise in datasets like 16S rRNA gene sequencing and resting-state fMRI, further complicated by small sample sizes observed in similar studies ^21–23^. To overcome these limitations, we employed a systematic feature selection strategy, combining ANOVA for initial feature identification, random forest-based importance ranking for dimensionality reduction, and partial correlation analysis to mitigate multicollinearity ^24–26^.

This rigorous methodology successfully narrowed the feature space to the most biologically informative variables ^27,28^, enhancing interpretability and reducing the likelihood of false-positive associations ^29–31^. The integration of fecal and saliva 16S rRNA datasets with neuroimaging features enabled a comprehensive exploration of microbial diversity and its interactions with neural correlates of CI, aligning with prior multi-omics studies ^32–34^.

Our integrated model outperformed stand-alone models, with Random Forest achieving the highest validation accuracy (0.889) and AUC (0.972), highlighting the synergistic potential of combining multimodal data ^35,36^. These findings demonstrate the utility of advanced feature selection and machine learning techniques in identifying robust biomarkers and gut-brain interactions relevant to neurodegenerative diseases.

### 4.2 Superior Classification of Cognitive Impairment Stages

A key challenge in PD research is the differentiation of MCI from more advanced cognitive states. Prior studies integrating multi-omics data in PD have reported moderate performance in distinguishing MCI from PDD. The proposed model exhibited enhanced specificity in detecting MCI, indicating that microbiome-derived features contribute significant predictive value beyond neuroimaging-based assessments ^37,38^.

In comparative studies focusing on Parkinson’s disease, similar or slightly lower performance metrics have been reported. For example, one study utilizing multi-omics data integration reported an AUC of 0.93 and accuracy of 0.81 in predicting cognitive impairment. Notably, our model demonstrates improved differentiation of MCI from more advanced cognitive states compared to the findings of this study, which reported limitations in specificity and Matthews Correlation Coefficient (MCC) in distinguishing MCI from dementia (AUC 0.69–0.75, MCC 0.177–0.470) ^38^. This highlights the potential of our multi-modals framework for resolving intermediate cognitive stages with greater accuracy.

Another study achieved an AUC of 0.91 using only fMRI features ^39^. Additionally, research on Alzheimer’s disease reported a predictive accuracy of 0.883 and an AUC of 0.970 using a framework that integrates explainable boosting machines (EBMs) with deep learning-based feature extraction on neuroimaging data from the ADNI dataset ^40^. Moreover, compared to studies using only fMRI features or deep learning approaches in neurodegenerative disease classification, the findings here validate the enhanced predictive power of integrating gut and oral microbiota with neuroimaging markers ^11,39,40^. The ability to capture subtle cognitive changes at early disease stages highlights the clinical relevance of this approach and its potential for early intervention strategies.

### 4.3 Identifying Neural and Microbial Signatures of Cognitive Decline

This study identifies previously unreported associations between functional brain connectivity and microbial composition in PD. Alterations in the default mode, salience, and attention networks emerged as key indicators of cognitive impairment, consistent with prior research ^41–43^ linking these circuits to cognitive regulation. Connectivity disruptions in these networks have been associated with neurodegeneration, supporting their potential as imaging biomarkers for monitoring disease progression.

In the gut microbiome, beneficial taxa like *Faecalibacterium* and *Lachnospira* decreased with disease progression, while pro-inflammatory taxa such as *Bacteroides* increased, supporting the role of dysbiosis in neuroinflammation and cognitive decline ^44,45^. Similarly, changes in salivary microbiota, including increased *Streptococcus, Veillonella,* and *Gemella,* further reinforce the relevance of microbial shifts in PD progression ^45–47^. These findings align with the gut-brain axis model, suggesting that microbial composition influences neural function through immune and metabolic pathways ^48^. The increase in Bacteroides and decrease in beneficial taxa such as *Faecalibacterium* in PD patients is consistent with previous findings linking dysbiosis to neuroinflammation and cognitive decline ^49,50^.

Machine learning models also revealed independent significant associations between fMRI connectivity features and specific microbial populations. Positive correlations between *Prevotella* and neural connectivity suggest potential interactions between microbiota and neural networks and reflect the dynamic interaction of gut and brain functions ^51,52^, while negative correlations between *Streptococcus* and limbic connectivity indicate possible neuroinflammatory effects. These findings underscore the interplay between the microbiome and brain function.

### 4.4 The Gut-Brain and Oral-Brain Interactions in PD Pathophysiology

This study is among the first to demonstrate concurrent alterations in gut and oral microbiota alongside neuroimaging changes in PD. The enrichment of *Veillonella* and *Streptococcus* in PDD suggests a pro-inflammatory microbial shift that may contribute to neuroinflammation and disease progression ^53–55^.

Negative associations between oral bacteria such as *Streptococcus* and brain connectivity highlight the sensitivity of the limbic system to intrinsic physiological changes and likely reflect neuroinflammatory responses ^56,57^. In addition, reductions in *Faecalibacterium* have been associated with altered somatomotor network activity, suggesting a bidirectional effect of motor network disruptions and microbial dysregulation ^44,48^.

Associations between *Veillonella* abundance and attention networks highlight the interaction between stress, inflammation, and oral microbiota, potentially mediated through the role of the salience network in interoceptive regulation ^58^. These findings are consistent with the gut-brain axis model in which microbial changes can modulate cognitive and motor functions, further confirming their role in PD progression ^59,60^. While previous research has predominantly focused on the gut-brain axis, the findings of this study underscore the oral microbiome as an additional factor contributing to PD-related neurodegeneration. The integration of oral microbiota alongside gut microbiota provides a novel perspective on the pathophysiology of the disease and suggests that microbial composition may serve as a potential biomarker for cognitive decline in PD. This highlights the need for a more comprehensive investigation of host-microbiome interactions across multiple body sites to better understand their role in neurodegenerative processes.

### 4.5 Microbiome, Demographics, and Brain Connectivity in Cognitive Impairment in Parkinson’s Disease

This study reveals significant associations between microbiome composition, demographic factors, and neural connectivity in cognitive impairment. Salivary *Bacteroides* and *Veillonella* were negatively correlated with age, while *Lachnospira* abundance positively associated with education, highlighting the impact of lifestyle and age on the microbiome ^61,62^.

Cognitive decline was linked to higher abundance levels of salivary *Streptococcus* and *Veillonella* (higher CDRS), whereas Neisseria abundance correlated positively with cognitive performance (higher MMSE), suggesting microbial contributions to neuroinflammation and neurotransmitter pathways ^63–66^.

fMRI analyses revealed that increased connectivity within the salience and limbic networks was associated with cognitive decline, suggesting these regions as potential markers of neurodegeneration. In contrast, stronger connections between the frontal and salience networks were linked to better cognitive performance, highlighting their importance for maintaining healthy brain function ^67–70^. These findings suggest that targeting the microbiome could help influence brain function and potentially slow the progression of neurodegenerative diseases like Parkinson’s by modulating gut-brain interactions.

### 4.6 Study Limitations

This study underscores the potential of integrating neuroimaging and microbiome data to explore cognitive impairment in Parkinson’s disease. Despite its strengths, however, this study has several limitations. The sample size, while sufficient for exploratory analyses, requires validation in larger and more diverse cohorts. Additionally, potential confounders such as diet, medication, and comorbidities were not extensively analyzed but are known to impact both microbiome composition and neuroimaging measures. Future studies should incorporate these variables to refine observed associations.

The cross-sectional design limits causal inferences regarding microbiome-neuroimaging interactions. Longitudinal studies are needed to determine whether microbial changes precede or follow neurodegenerative alterations. Additionally, the exclusion of PD patients without cognitive impairment may confound the findings in predicting CI in PD.

Despite these limitations, our study provides a strong foundation for future research by identifying potential biomarkers and elucidating novel microbiota-brain interactions. These insights contribute to an increasing understanding of the gut-brain axis and support the development of microbiome-targeted interventions for neurodegenerative diseases.

## 5 CONCLUSIONS

This study highlights the value of integrating neuroimaging and microbiome data to provide insights into complex interactions between microbial dysbiosis, neural connectivity, and cognitive impairment in Parkinson’s disease. By delineating major microbial and neural biomarkers and their stage-dependent interrelations, findings provide firm bases for the development of stage specific, targeted therapeutic strategies that are aimed at modulating the gut-brain axis. Subsequent studies should be focused on more diverse and larger cohorts and examine mechanisms by which microbiota modulate neural function to advance precision medicine strategies in PD and other neurodegenerative disease.

## AUTHOR CONTRIBUTIONS

SY conceived and designed the study and supervised the research. BD developed and implemented the analytical methods, performed data analysis, and prepared the initial manuscript draft. UN provided intellectual contributions to method development and interpretation of findings. All authors contributed to writing and revising the manuscript and approved the final version

## FUNDING

This study was supported by a grant to Suleyman Yildirim from the Scientific and Technological Research Council of Turkey (TÜBITAK) (grant no. 315S301)

## CONFLICT OF INTEREST

The authors confirm that this research was conducted independently and without any commercial or financial interests that could be perceived as potential conflicts of interest.

## DATA AVAILABILITY STATEMENT

The data that support the findings of this study are openly available in the NCBI Bio project at Stool: https://www.ncbi.nlm.nih.gov/bioproject/PRJNA1184470, Saliva: https://www.ncbi.nlm.nih.gov/bioproject/PRJNA1184768.

## SUPPORTING INFORMATION

S1. Supporting Information File.

S2. Supporting Information File. Materials and Methods

S3. Supplementary Table 1-Functional Network Connectivity Feature Names Supplementary Table 2- Functional Network Connectivity fMRI Final Selected Features

## Notes

### Competing Interest Statement

The authors have declared no competing interest.

