## Supplementary material for "Integrative Analysis of Neuroimaging and Microbiome Data Predicts Cognitive Decline in Parkinson’s Disease": S1. Supporting Information File. S2. Supporting Information File. Materials and Methods S3. Supplementary Table: S1_figures_ML_article.docx

**Table S1.1.** Hyperparameters optimization of fMRI dataset independent models for feature selection and machine learning classification algorithms of study groups.

| Model | parameters |
| --- | --- |
| Feature selection (RF) | n estimators=100. criterion= Gini Impurity. max features=15 |
| Random Forest | n estimators=100. criterion= Gini Impurity  Best Hyperparameters: {'bootstrap': True. 'max_depth': 10. 'min_samples_leaf': 2. 'min_samples_split': 10. 'n_estimators': 200} |
| DT | Best Hyperparameters: {'criterion': 'entropy'. 'max_depth': 10. 'max_features': None. 'min_samples_leaf': 5. 'min_samples_split': 5. 'splitter': 'best'} |
| SVM | Best Hyperparameters: {'C': 125. 'gamma': 1. 'kernel': 'linear'} |
| Adaboost | Best Hyperparameters: {'learning_rate': 0.4. 'n_estimators': 110} |
| XGboost | Best Hyperparameters: {'learning_rate': 0.01. 'n_estimators': 50} |
| Logistic regression | Best Hyperparameters: {'C': 100. 'penalty': 'l1'. 'solver': 'liblinear'} |

**Table S1.2.** Hyperparameters optimization of Stool ASV dataset independent models for feature selection and machine learning classification algorithms of study groups.

| Model | parameters |
| --- | --- |
| Feature selection (Lasso) | Best alpha parameter: 0.08087 |
| Random Forest | Best Hyperparameters: {'bootstrap': False. 'max_depth': 25. 'min_samples_leaf': 1. 'min_samples_split': 5. 'n_estimators': 100} |
| DT | Best Hyperparameters: {'criterion': 'gini'. 'max_depth': 20. 'max_features': 'sqrt'. 'min_samples_leaf': 2. 'min_samples_split': 5. 'splitter': 'best'} |
| SVM | Best Hyperparameters: {'C': 0.5. 'gamma': 0.001. 'kernel': 'rbf'} |
| Adaboost | Best Hyperparameters: {'learning_rate': 0.01. 'n_estimators': 50} |
| XGboost | Best Hyperparameters: {'learning_rate': 0.01. 'n_estimators': 50} |
| Logistic regression | Best Hyperparameters: {'C': 80. 'penalty': 'none'.'solver': 'liblinear'} |

**Table S1.3.** Hyperparameters optimization of Saliva ASV dataset independent models for feature selection and machine learning classification algorithms of study groups.

| Model | parameters |
| --- | --- |
| Feature selection (Lasso) | Best alpha parameter: 0.07034 |
| Random Forest | Best Hyperparameters: {'bootstrap': True. 'max_depth': 25. 'min_samples_leaf': 2. 'min_samples_split': 5. 'n_estimators': 200} |
| DT | Best Hyperparameters: {'criterion': 'gini'. 'max_depth': 20. 'max_features': 'sqrt'. 'min_samples_leaf': 2. 'min_samples_split': 5. 'splitter': 'best'} |
| SVM | Best Hyperparameters: {'C': 10. 'gamma': 0.001. 'kernel': 'rbf'} |
| Adaboost | Best Hyperparameters: {'learning_rate': 0.5. 'n_estimators': 50} |
| XGboost | Best Hyperparameters: {'learning_rate': 0.4. 'n_estimators': 200} |
| Logistic regression | Best Hyperparameters: {'C': 0.01. 'penalty': 'none'. 'solver': 'saga'} |

**Table S1.4.** Hyperparameters optimization of joint model (fMRI+Stool+Saliva ASV datasets) for feature selection and machine learning classification algorithms of study groups.

**all random state number=42*

| Model | parameters |
| --- | --- |
| Feature selection (Lasso) | Best alpha parameter: 0.08253 |
| Random Forest | Best Hyperparameters: {'bootstrap': True. 'max_depth': 10. 'min_samples_leaf': 1. 'min_samples_split': 10. 'n_estimators': 300} |
| DT | Best Hyperparameters: {'criterion': 'gini'. 'max_depth': 10. 'max_features': None. 'min_samples_leaf': 5. 'min_samples_split': 5. 'splitter': 'random'} |
| SVM | Best Hyperparameters: {'C': 0.1. 'gamma': 1. 'kernel': 'linear'} |
| Adaboost | Best Hyperparameters: {'learning_rate': 0.2. 'n_estimators': 100} |
| XGboost | Best Hyperparameters: {'learning_rate': 0.4. 'n_estimators': 100} |
| Logistic regression | Best Hyperparameters: {'C': 100. 'penalty': 'l1'. 'solver': 'liblinear} |

**Table S2.1.** Performance metric scores of ML classification algorithms for the rs-fMRI independent model.

|  | Training dataset (n = 42) | | | | | Validation dataset (n = 18) | | | | |
| --- | --- | --- | --- | --- | --- | --- | --- | --- | --- | --- |
| Model | Accuracy | Precision | Recall | F1 Score | AUC Score | Accuracy | Precision | Recall | F1 Score | AUC Score |
| Random Forest (RF) | 0.905 | 0.926 | 0.904 | 0.907 | 0.992 | 0.833 | 0.849 | 0.833 | 0.837 | 0.944 |
| Support Vector Machine (SVM) | 0.952 | 0.958 | 0.956 | 0.954 | 0.958 | 0.5 | 0.525 | 0.5 | 0.506 | 0.764 |
| Decision tree (DT) | 0.857 | 0.865 | 0.857 | 0.858 | 0.969 | 0.611 | 0.611 | 0.611 | 0.602 | 0.59 |
| AdaBoost | 0.929 | 0.933 | 0.93 | 0.931 | 0.992 | 0.667 | 0.679 | 0.667 | 0.662 | 0.778 |
| XGBoost | 1.0 | 1.0 | 1.0 | 1.0 | 1.0 | 0.556 | 0.581 | 0.556 | 0.554 | 0.806 |
| Logistic regression | 0.952 | 0.954 | 0.954 | 0.954 | 0.983 | 0.556 | 0.565 | 0.556 | 0.558 | 0.815 |

**Figure S1.1.** ROC curves of ML classification algorithms for the rs-fMRI independent model.

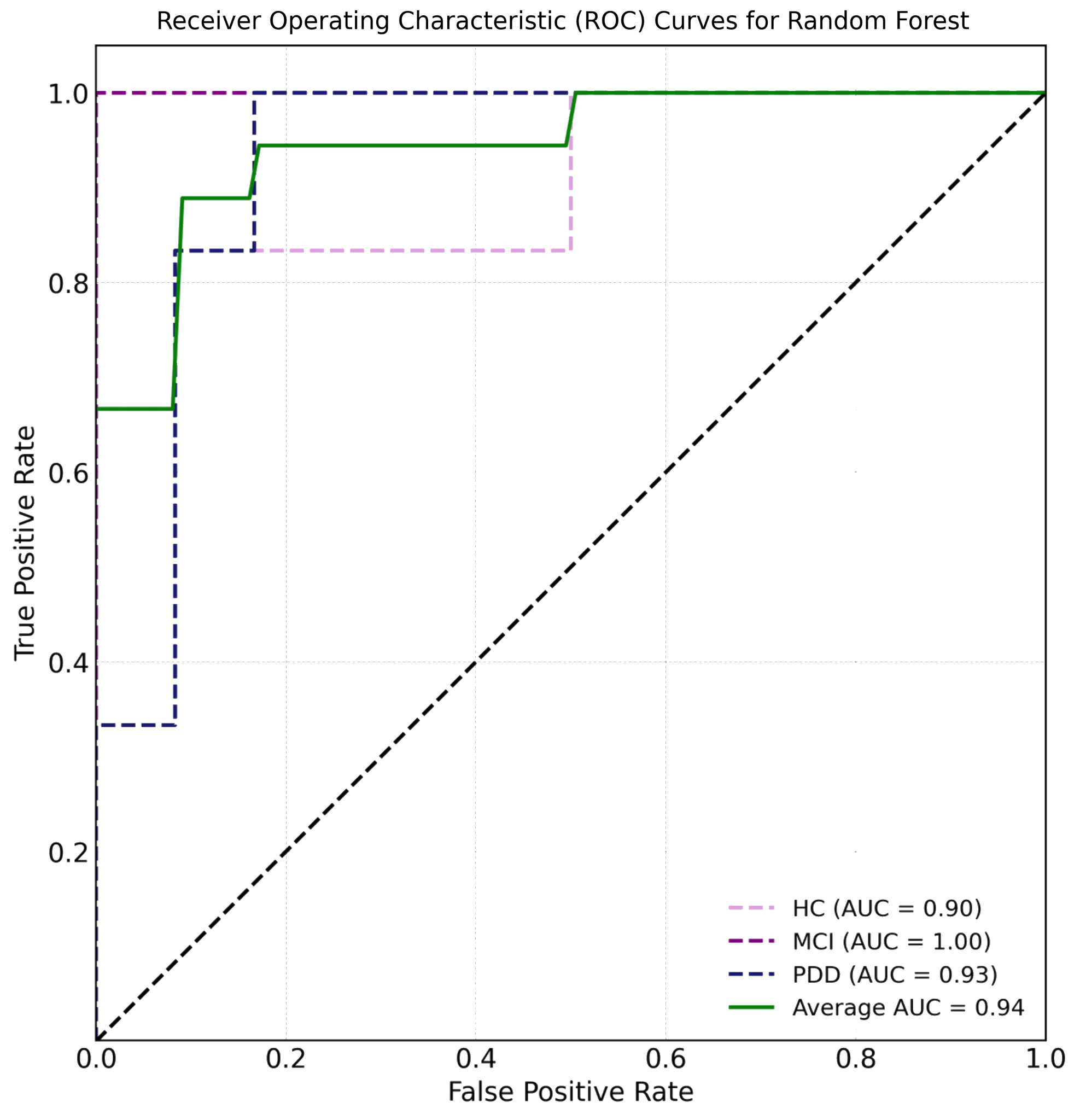

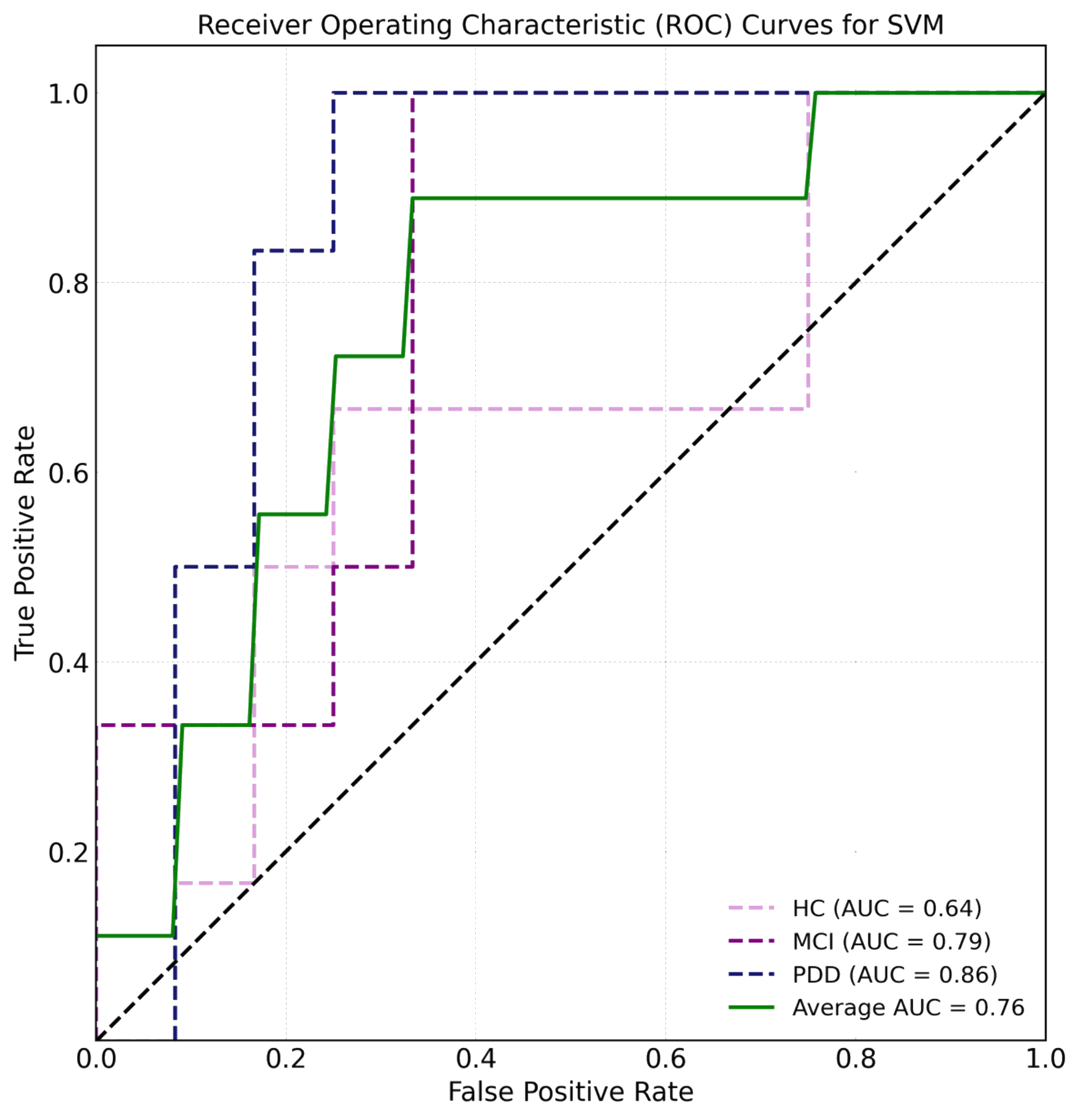

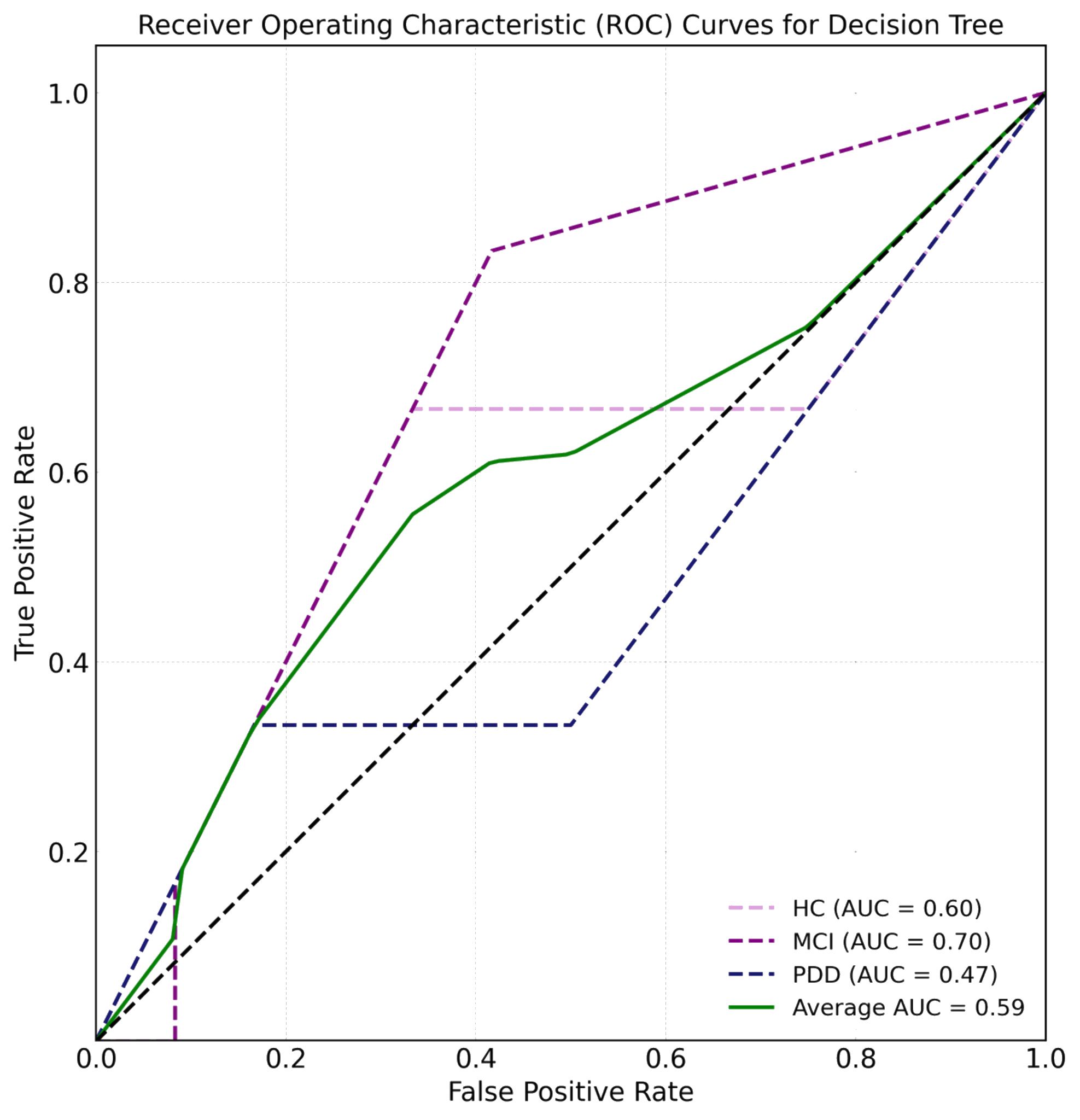

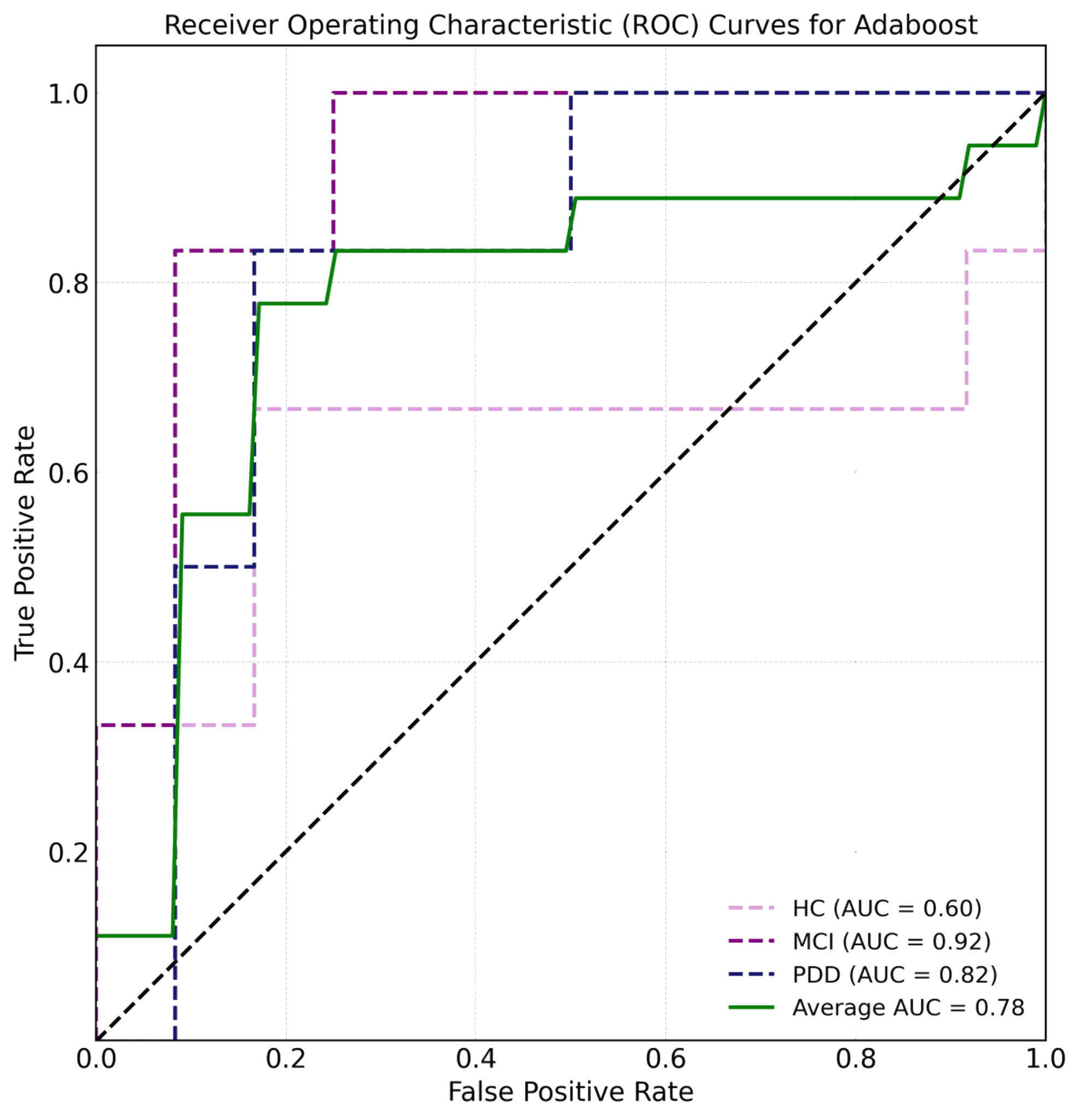

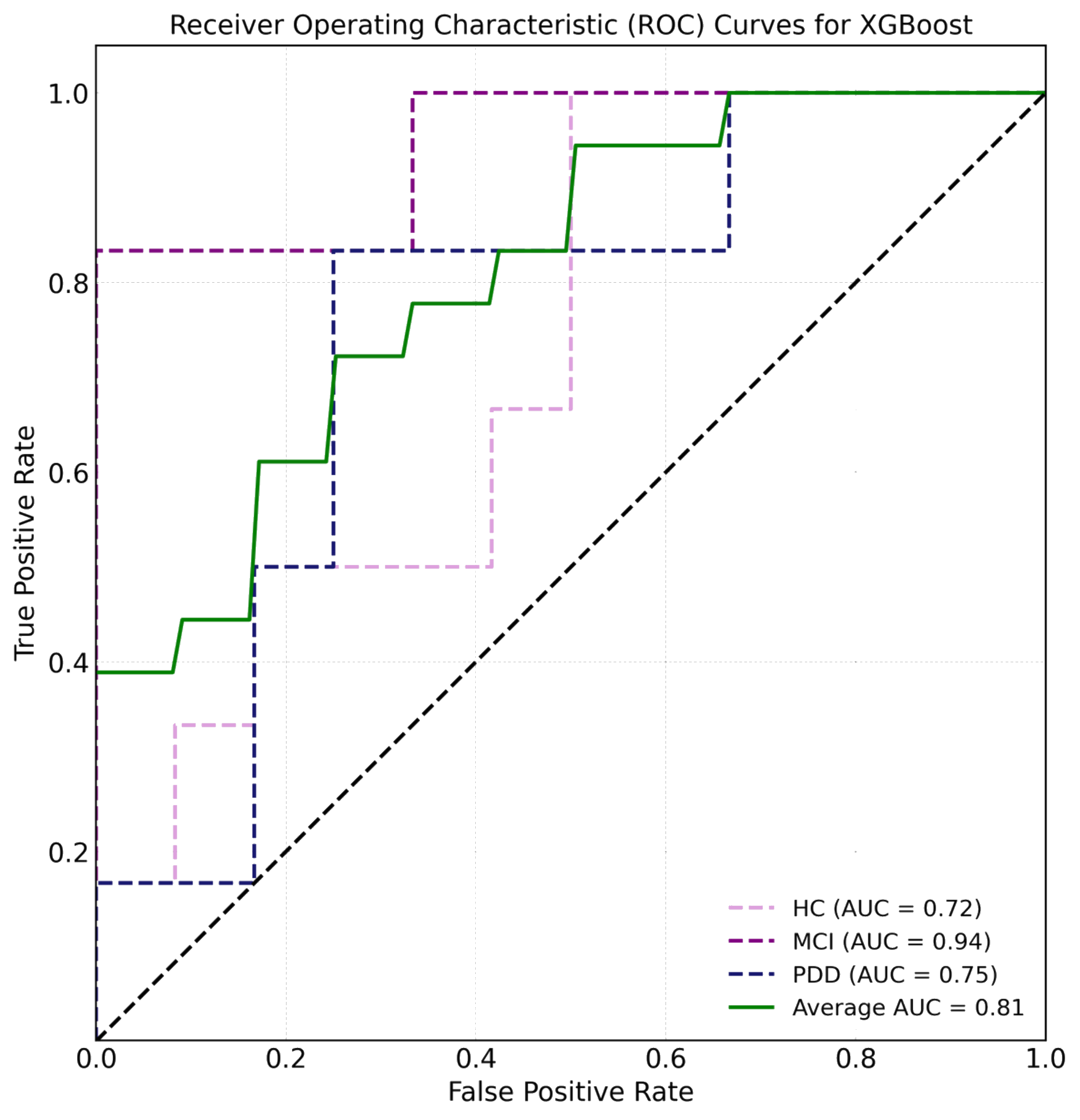

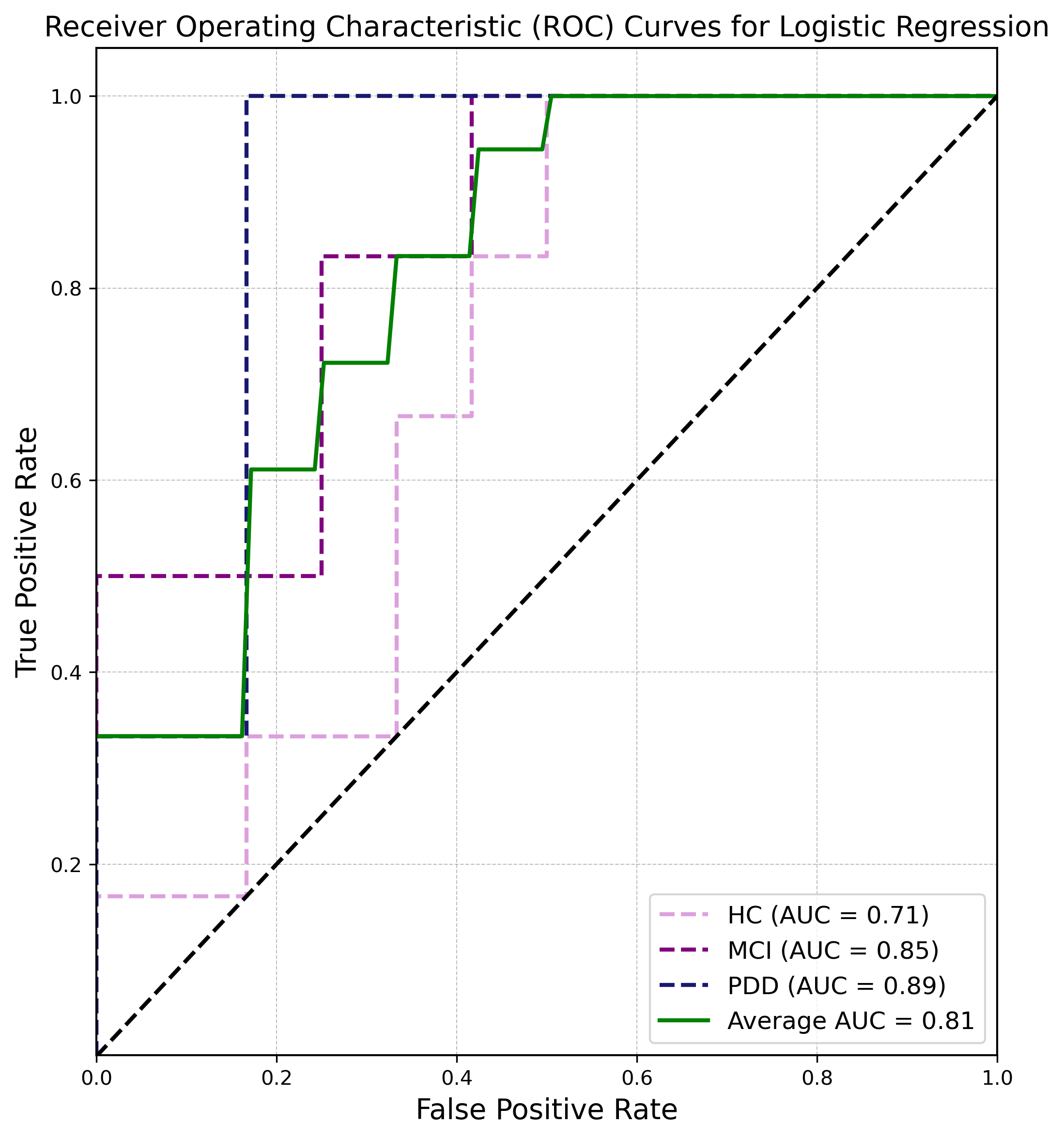

**Table S2.2.** Performance metric scores of ML classification algorithms for the Stool ASV dataset independent model.

|  | Training dataset (n = 42) | | | | | Validation dataset (n = 18) | | | | |
| --- | --- | --- | --- | --- | --- | --- | --- | --- | --- | --- |
| Model | Accuracy | Precision | Recall | F1 Score | AUC Score | Accuracy | Precision | Recall | F1 Score | AUC Score |
| Random Forest (RF) | 0.786 | 0.856 | 0.782 | 0.779 | 0.937 | 0.611 | 0.556 | 0.611 | 0.563 | 0.794 |
| Support Vector Machine (SVM) | 0.524 | 0.352 | 0.526 | 0.422 | 0.7 | 0.667 | 0.45 | 0.667 | 0.536 | 0.75 |
| Decision tree (DT) | 0.667 | 0.706 | 0.67 | 0.674 | 0.848 | 0.5 | 0.556 | 0.5 | 0.469 | 0.641 |
| AdaBoost | 0.595 | 0.672 | 0.584 | 0.561 | 0.831 | 0.556 | 0.667 | 0.556 | 0.522 | 0.736 |
| XGBoost | 0.643 | 0.722 | 0.646 | 0.647 | 0.836 | 0.444 | 0.675 | 0.444 | 0.41 | 0.639 |
| Logistic regression | 1.0 | 1.0 | 1.0 | 1.0 | 1.0 | 0.833 | 0.838 | 0.833 | 0.832 | 0.782 |

**Figure S1.2.** ROC curves of ML classification algorithms for the Stool ASV dataset independent model.

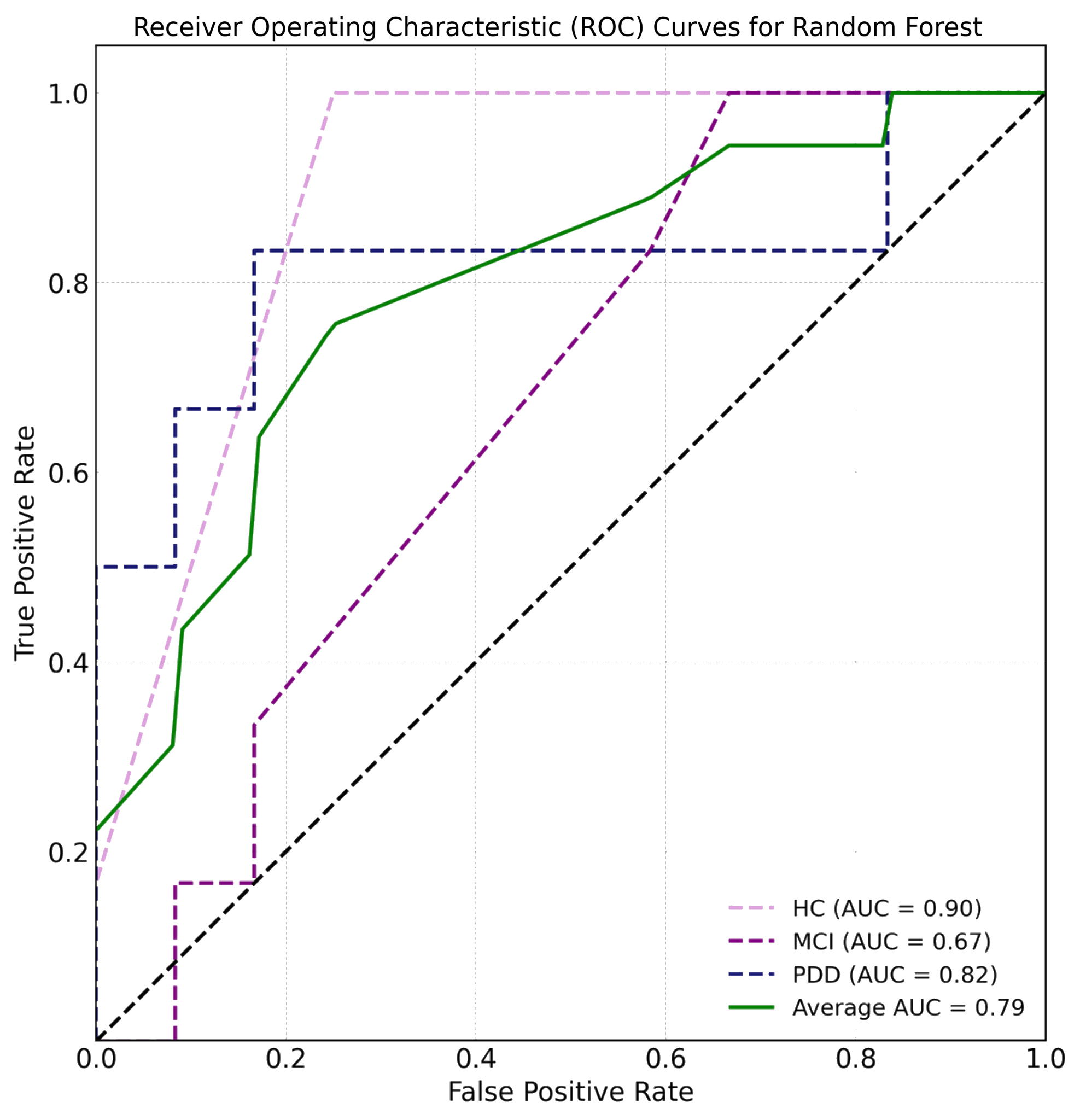

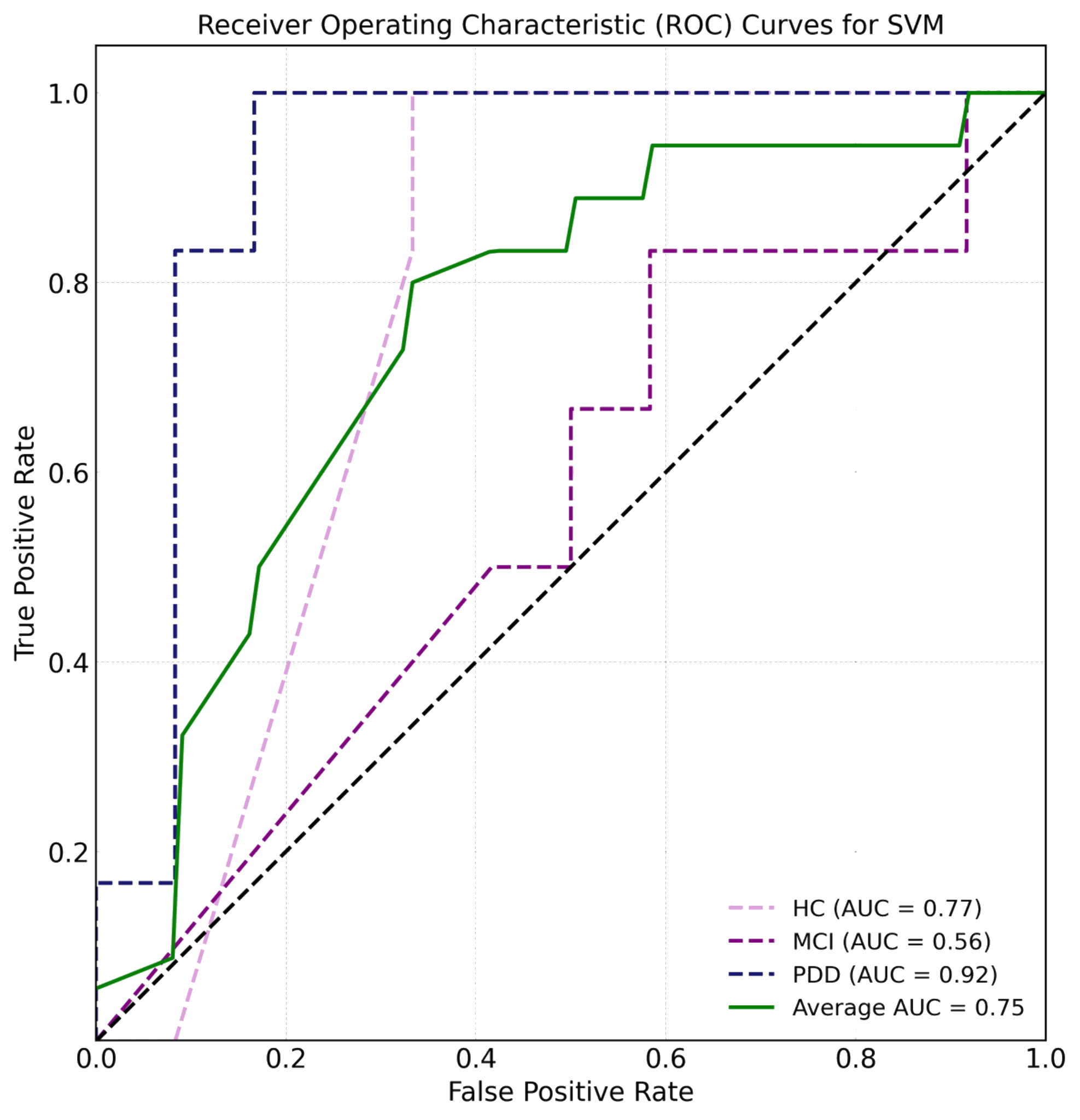

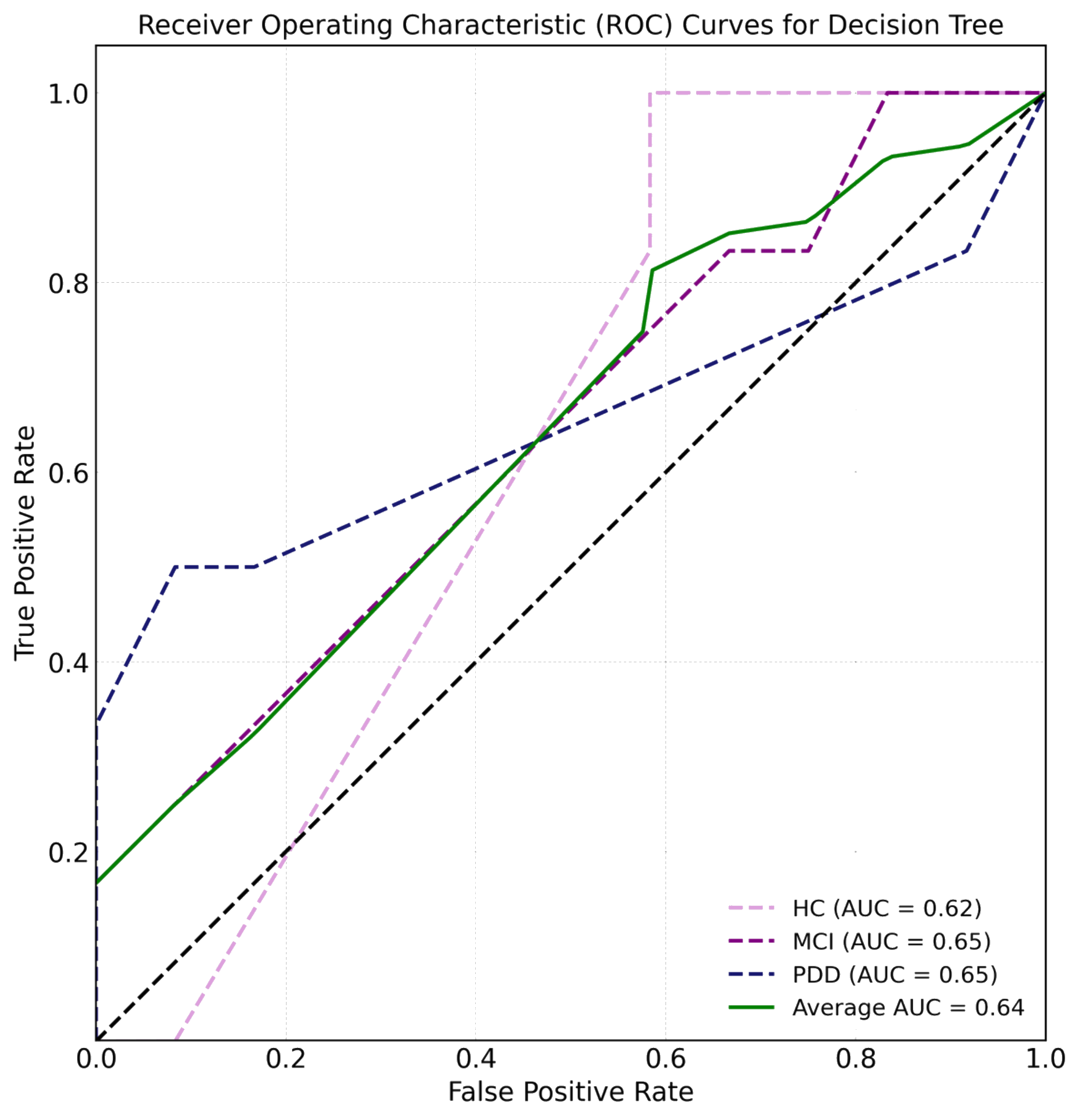

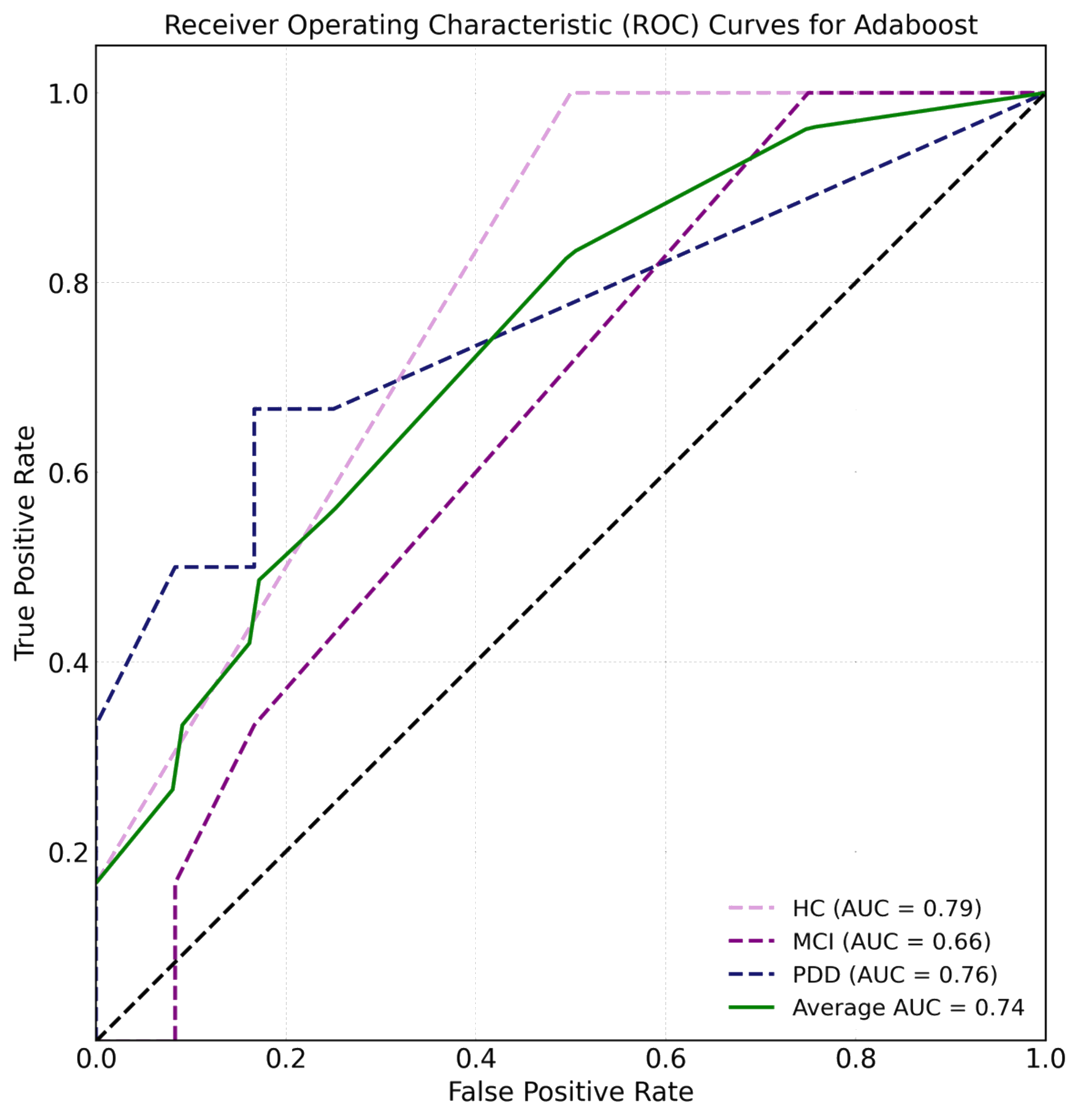

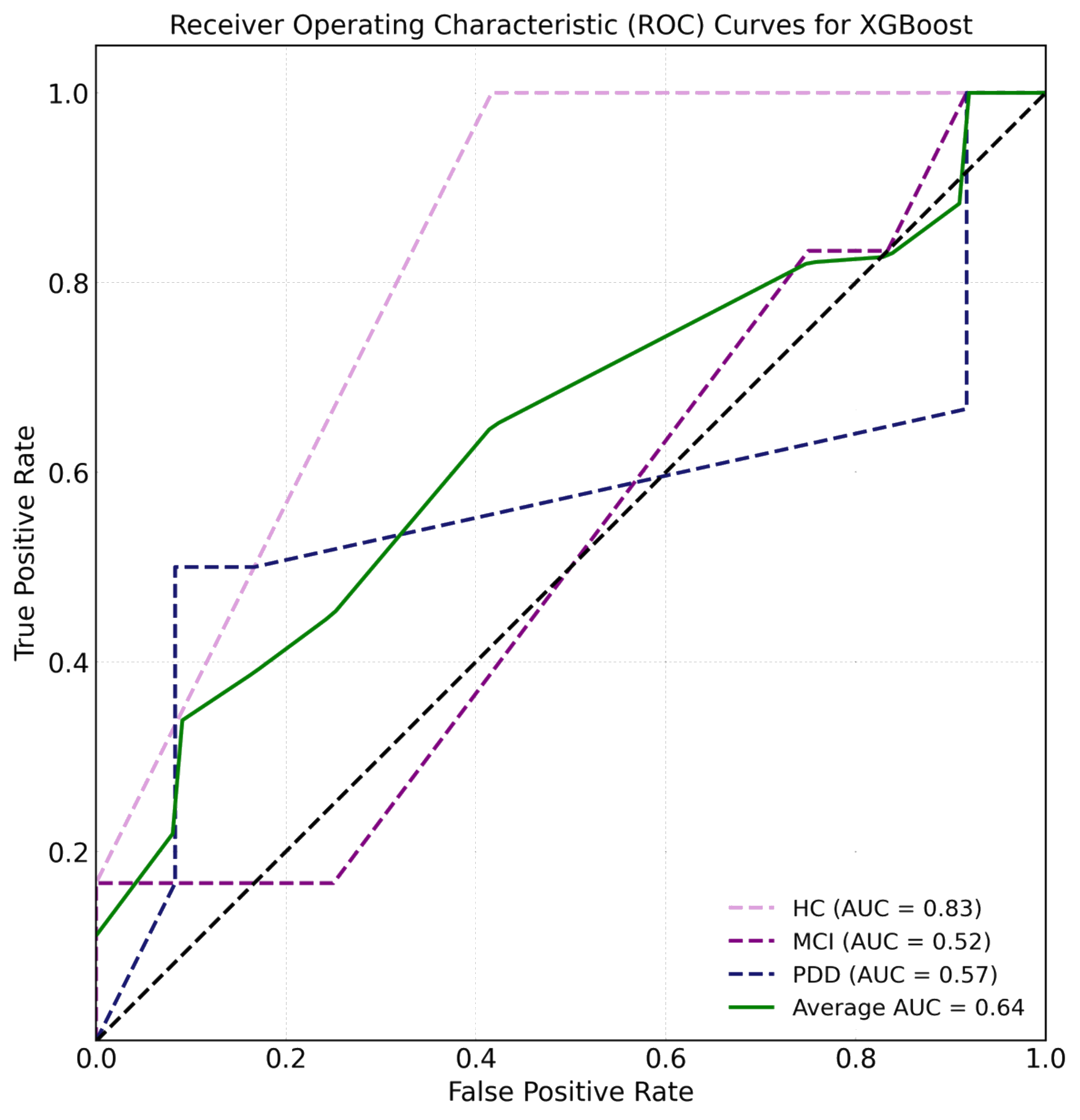

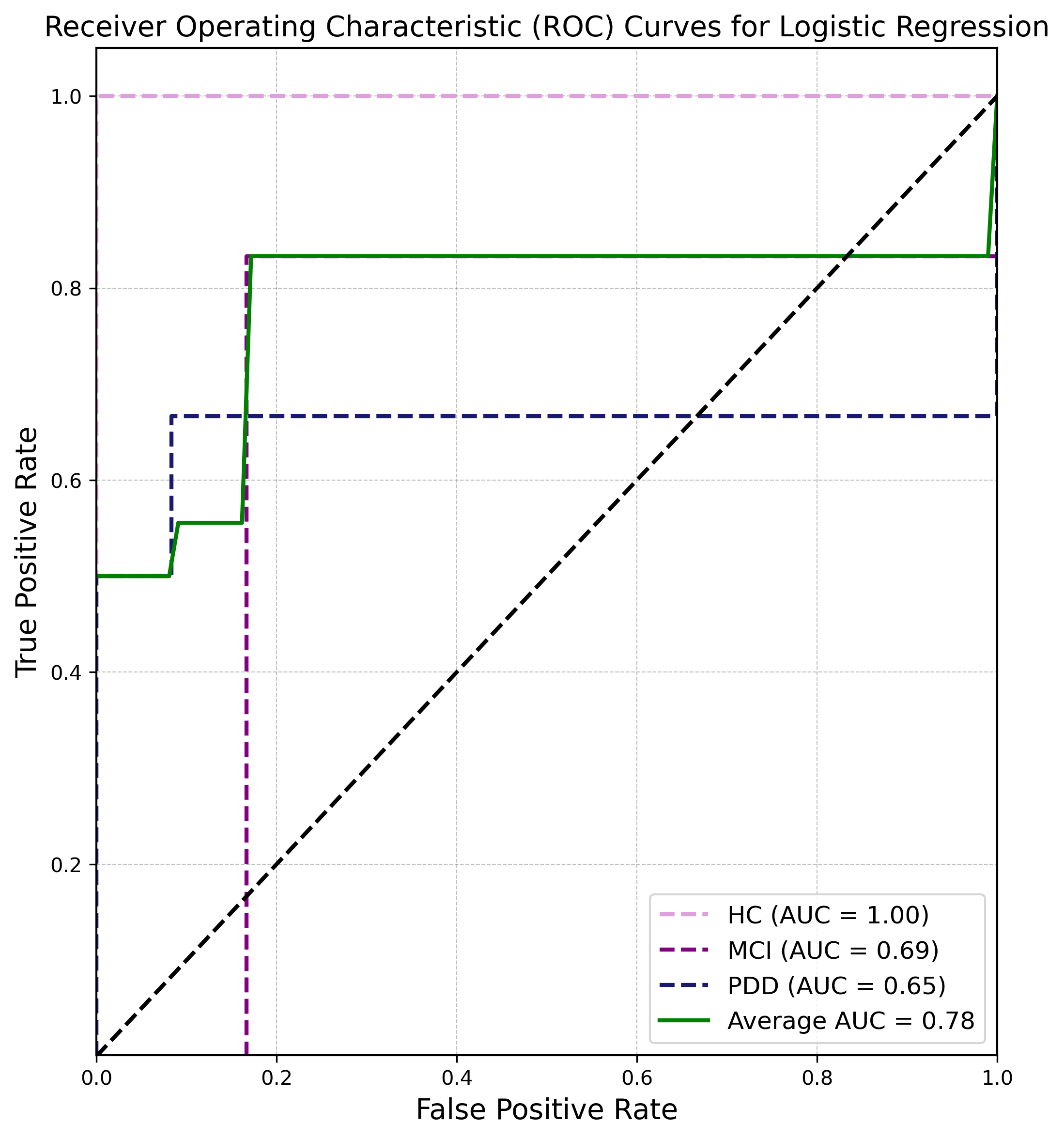

**Table S2.3.** Performance metric scores of ML classification algorithms for the Saliva ASV dataset independent model.

|  | Training dataset (n = 42) | | | | | Validation dataset (n = 18) | | | | |
| --- | --- | --- | --- | --- | --- | --- | --- | --- | --- | --- |
| Model | Accuracy | Precision | Recall | F1 Score | AUC Score | Accuracy | Precision | Recall | F1 Score | AUC Score |
| Random Forest (RF) | 1.0 | 1.0 | 1.0 | 1.0 | 0.998 | 0.556 | 0.611 | 0.556 | 0.519 | 0.736 |
| Support Vector Machine (SVM) | 1.0 | 1.0 | 1.0 | 1.0 | 1.0 | 0.444 | 0.452 | 0.444 | 0.441 | 0.639 |
| Decision tree (DT) | 0.738 | 0.744 | 0.738 | 0.727 | 0.897 | 0.444 | 0.444 | 0.444 | 0.444 | 0.63 |
| AdaBoost | 0.857 | 0.888 | 0.853 | 0.854 | 0.982 | 0.5 | 0.522 | 0.5 | 0.472 | 0.722 |
| XGBoost | 1.0 | 1.0 | 1.0 | 1.0 | 1.0 | 0.611 | 0.654 | 0.611 | 0.583 | 0.741 |
| Logistic regression | 0.857 | 0.862 | 0.863 | 0.856 | 0.978 | 0.5 | 0.633 | 0.5 | 0.5 | 0.745 |

**Figure S1.3.** ROC curves of ML classification algorithms for the Saliva ASV dataset independent model.

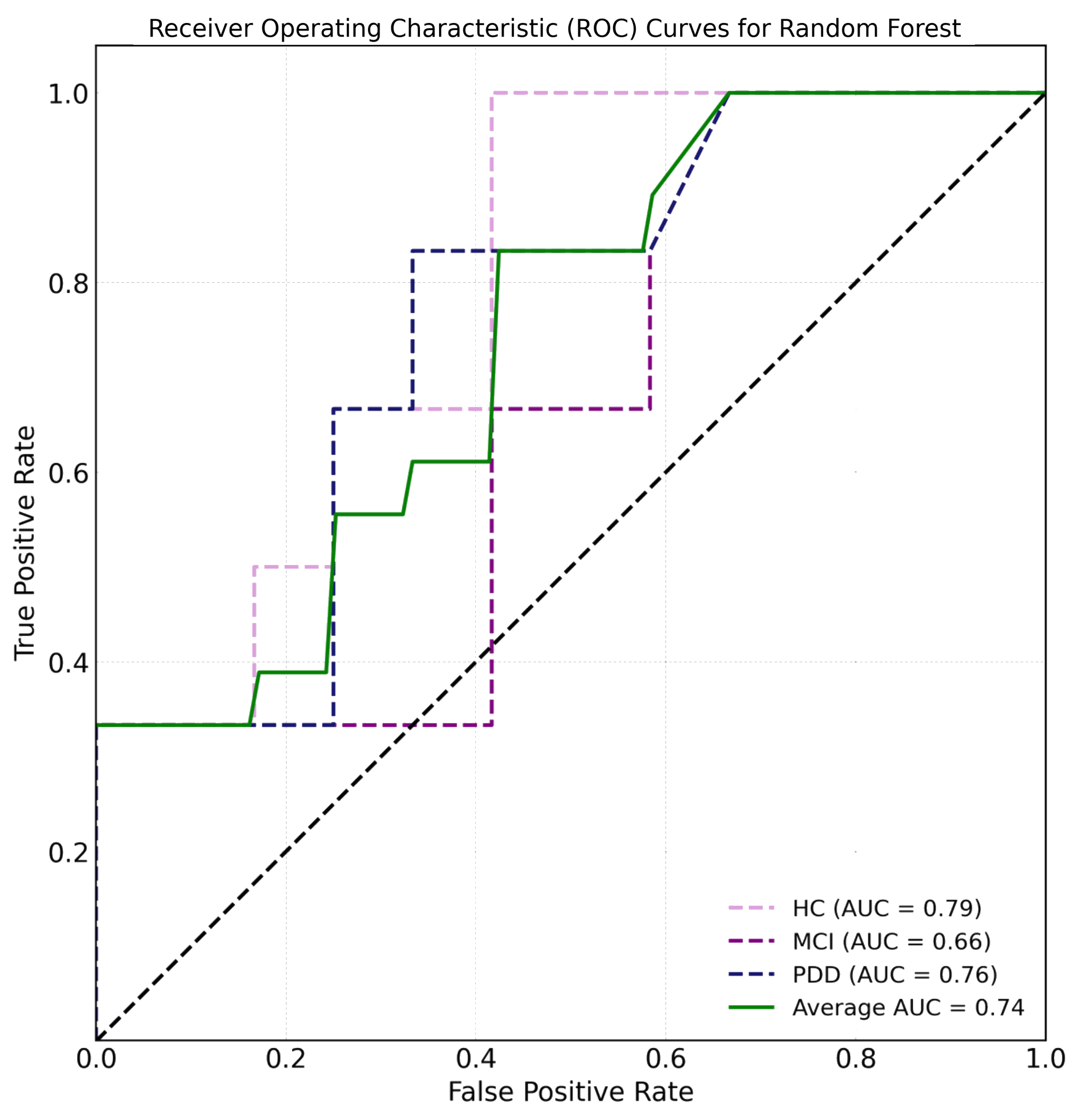

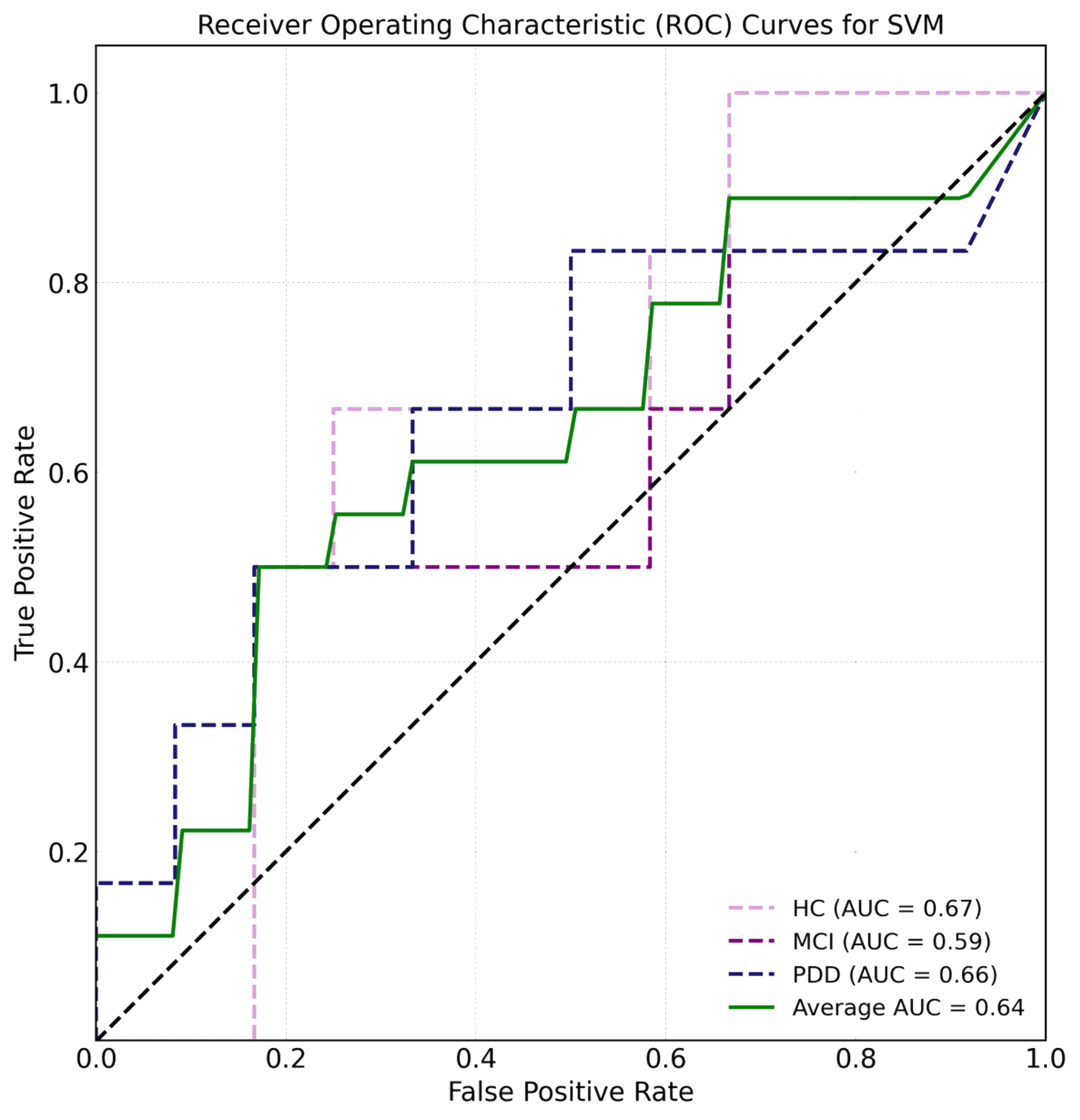

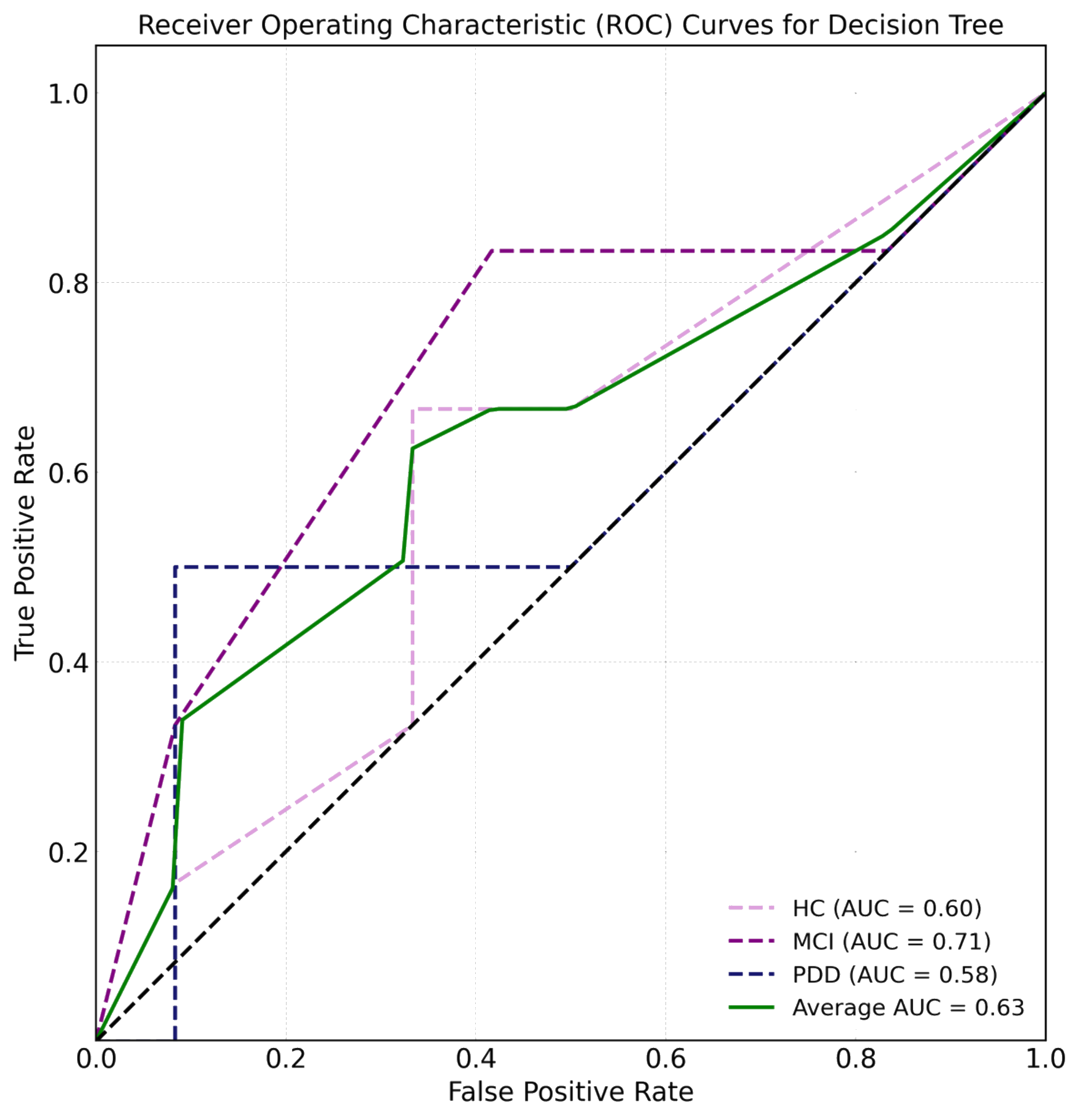

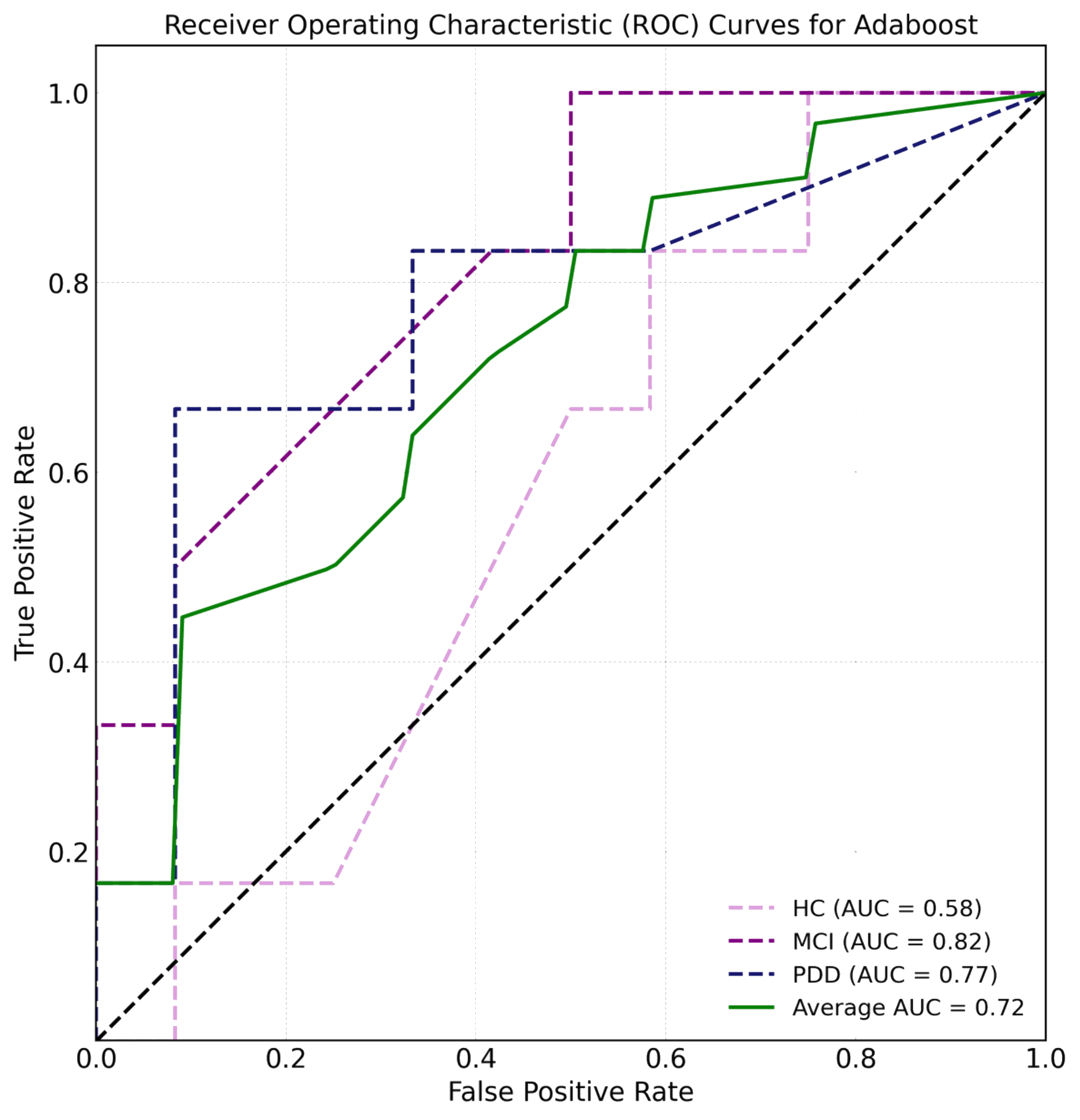

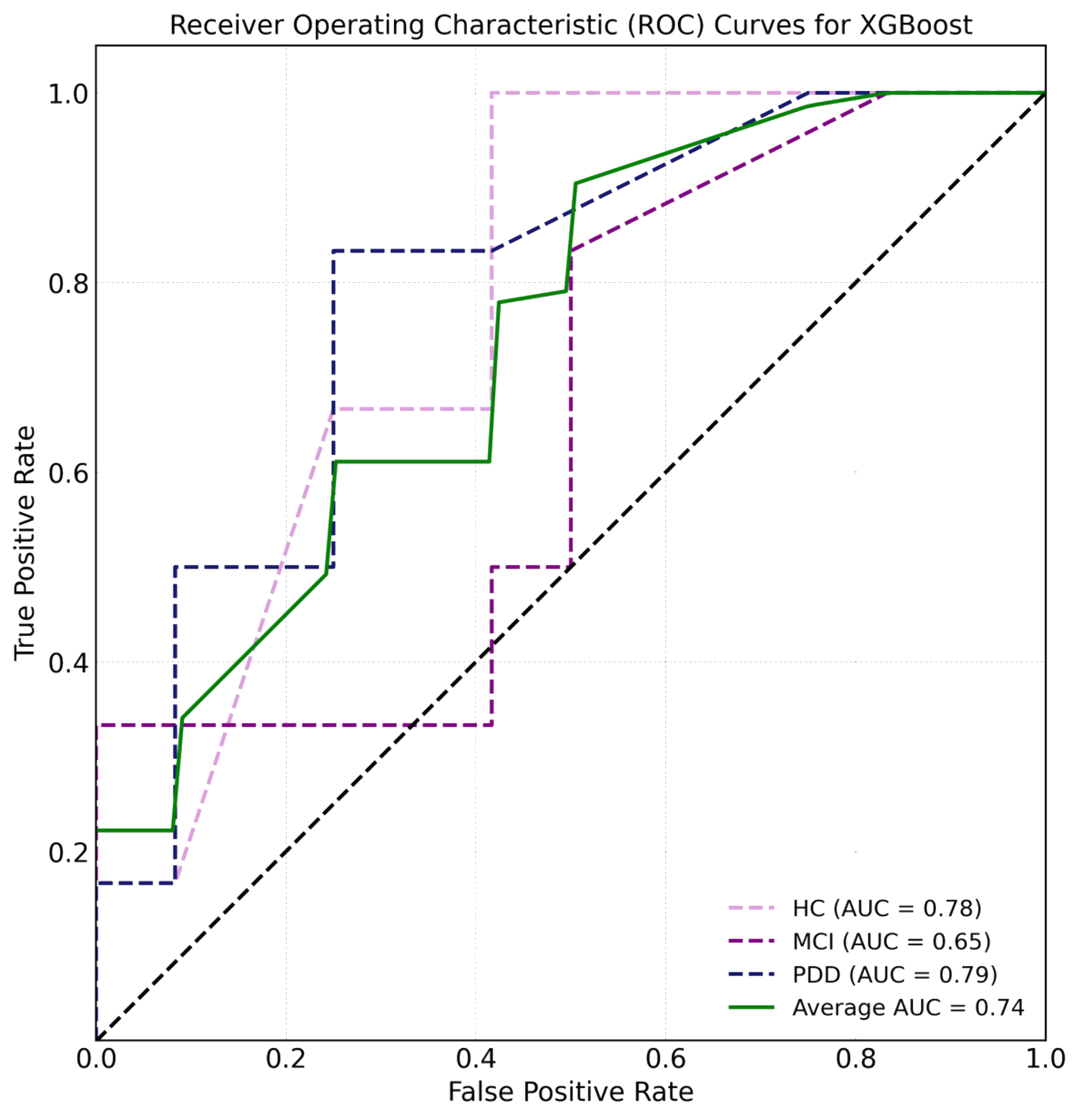

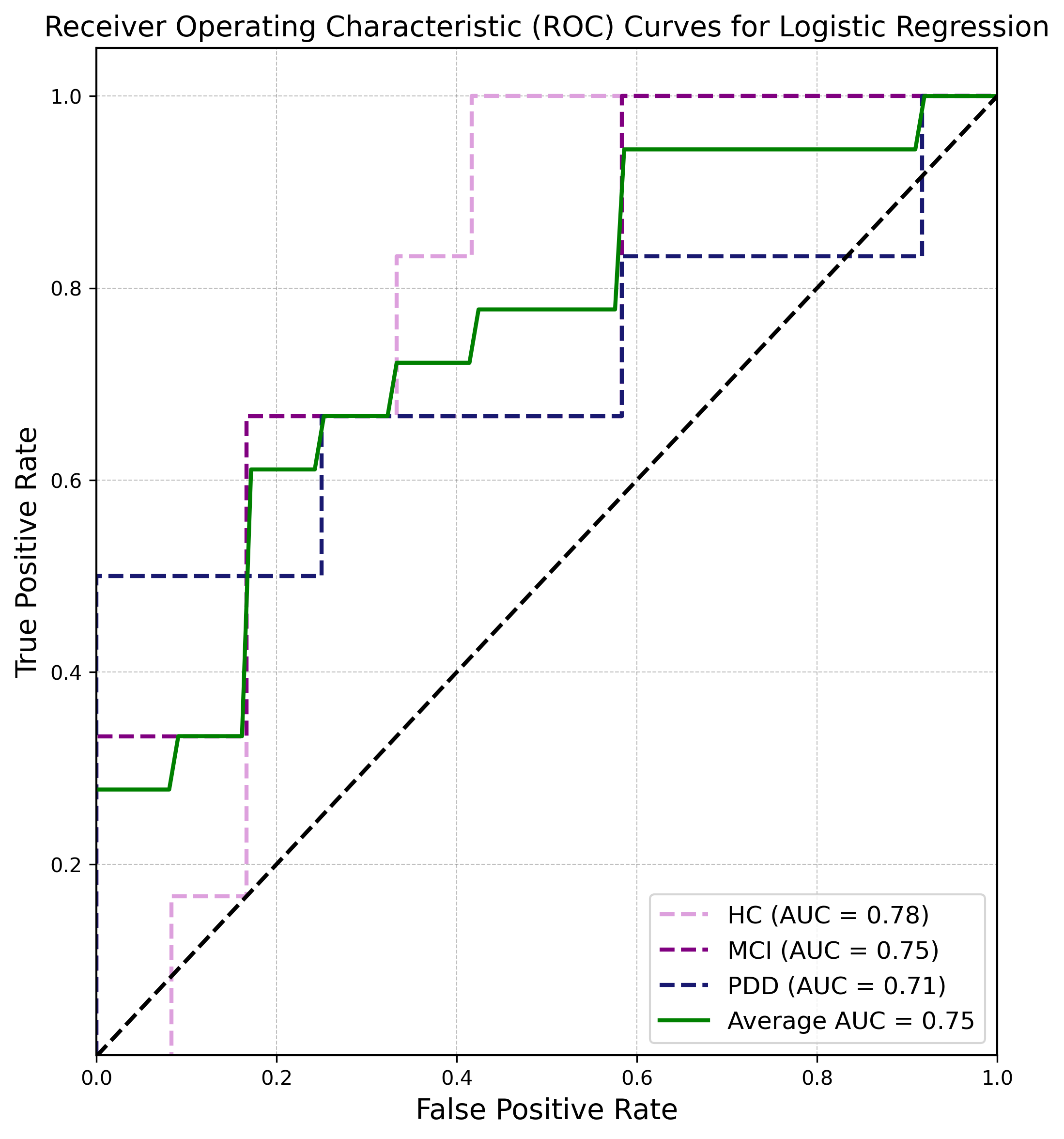

**Table S2.4.** Comparison of performance measurement scores of ML classification algorithms for independent models and joint model (fMRI+Feces+Saliva ASV datasets)

| Performance Metrics | Random Forest (RF) | Support Vector Machine (SVM) | Decision Tree (DT) | AdaBoost | XGBoost | Logistic Regression |
| --- | --- | --- | --- | --- | --- | --- |
| Accuracy_fMRI | 0.833 | 0.5 | 0.611 | 0.667 | 0.556 | 0.556 |
| Precision_fMRI | 0.849 | 0.525 | 0.611 | 0.679 | 0.581 | 0.565 |
| Recall_fMRI | 0.833 | 0.5 | 0.611 | 0.667 | 0.556 | 0.556 |
| F1 Score_fMRI | 0.837 | 0.506 | 0.602 | 0.662 | 0.554 | 0.558 |
| AUC Score_fMRI | 0.944 | 0.764 | 0.59 | 0.778 | 0.806 | 0.815 |
| Accuracy_Stool | 0.611 | 0.667 | 0.5 | 0.556 | 0.444 | 0.833 |
| Precision_Stool | 0.556 | 0.45 | 0.556 | 0.667 | 0.675 | 0.838 |
| Recall_Stool | 0.611 | 0.667 | 0.5 | 0.556 | 0.444 | 0.833 |
| F1 Score_Stool | 0.563 | 0.536 | 0.469 | 0.522 | 0.41 | 0.832 |
| AUC Score_Stool | 0.794 | 0.75 | 0.641 | 0.736 | 0.639 | 0.782 |
| Accuracy_Saliva | 0.556 | 0.444 | 0.444 | 0.5 | 0.611 | 0.5 |
| Precision_Saliva | 0.611 | 0.452 | 0.444 | 0.522 | 0.654 | 0.633 |
| Recall_Saliva | 0.556 | 0.444 | 0.444 | 0.5 | 0.611 | 0.5 |
| F1 Score_Saliva | 0.519 | 0.441 | 0.444 | 0.472 | 0.583 | 0.5 |
| AUC Score_Saliva | 0.736 | 0.639 | 0.63 | 0.722 | 0.741 | 0.745 |
| Accuracy_joint model | 0.889 | 0.778 | 0.667 | 0.611 | 0.722 | 0.778 |
| Precision_joint model | 0.889 | 0.819 | 0.643 | 0.621 | 0.722 | 0.822 |
| Recall_joint model | 0.889 | 0.778 | 0.667 | 0.611 | 0.722 | 0.778 |
| F1 Score_joint model | 0.889 | 0.783 | 0.646 | 0.591 | 0.722 | 0.776 |
| AUC Score_joint model | 0.972 | 0.866 | 0.84 | 0.838 | 0.921 | 0.852 |

**Table S3.1.** Comparison of Joint Model AUC Scores Using DeLong's Test.

| **Model 1** | **Model 2** | **AUC 1** | **AUC 2** | **Variance 1** | **Variance 2** | **Z-Score** | **p**  **Value** |
| --- | --- | --- | --- | --- | --- | --- | --- |
| Random Forest (RF) | Support Vector Machine (SVM) | 0.9722 | 0.8657 | 0.1534 | 0.2129 | 0.1759 | 0.8603 |
| Random Forest (RF) | Decision tree (DT) | 0.9722 | 0.8403 | 0.1534 | 0.2284 | 0.2135 | 0.8309 |
| Random Forest (RF) | AdaBoost | 0.9722 | 0.838 | 0.1534 | 0.1578 | 0.2407 | 0.8098 |
| Random Forest (RF) | XGBoost | 0.9722 | 0.9213 | 0.1534 | 0.1432 | 0.0935 | 0.9255 |
| Random Forest (RF) | Logistic regression | 0.9722 | 0.852 | 0.1534 | 0.105 | 0.2365 | 0.813 |
| Support Vector Machine (SVM) | Decision tree (DT) | 0.8657 | 0.8403 | 0.2129 | 0.2284 | 0.0383 | 0.9694 |
| Support Vector Machine (SVM) | AdaBoost | 0.8657 | 0.838 | 0.2129 | 0.1578 | 0.0456 | 0.9636 |
| Support Vector Machine (SVM) | XGBoost | 0.8657 | 0.9213 | 0.2129 | 0.1432 | 0.0931 | 0.9258 |
| Support Vector Machine (SVM) | Logistic regression | 0.8657 | 0.852 | 0.2129 | 0.105 | 0.0244 | 0.9806 |
| Decision tree (DT) | AdaBoost | 0.8403 | 0.838 | 0.2284 | 0.1578 | 0.0037 | 0.997 |
| Decision tree (DT) | XGBoost | 0.8403 | 0.9213 | 0.2284 | 0.1432 | 0.1329 | 0.8943 |
| Decision tree (DT) | Logistic regression | 0.8403 | 0.852 | 0.2284 | 0.105 | 0.0203 | 0.9838 |
| AdaBoost | XGBoost | 0.838 | 0.9213 | 0.1578 | 0.1432 | 0.1519 | 0.8793 |
| AdaBoost | Logistic regression | 0.838 | 0.852 | 0.1578 | 0.105 | 0.0274 | 0.9782 |
| XGBoost | Logistic regression | 0.9213 | 0.852 | 0.1432 | 0.105 | 0.1391 | 0.8894 |

**Figure S2.1.** Comparison of Precision and F1-Score performance metrics between groups in independent models.

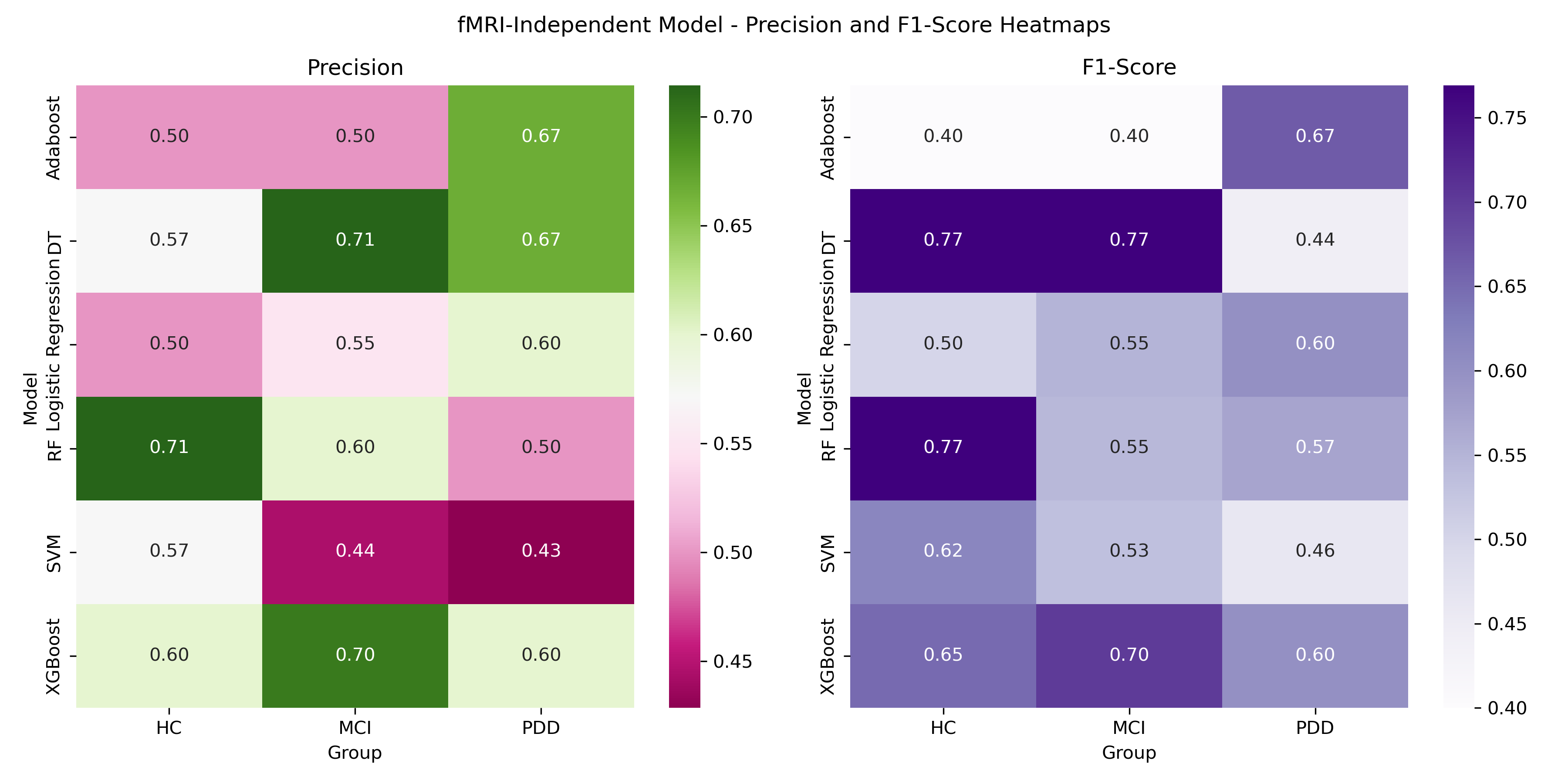

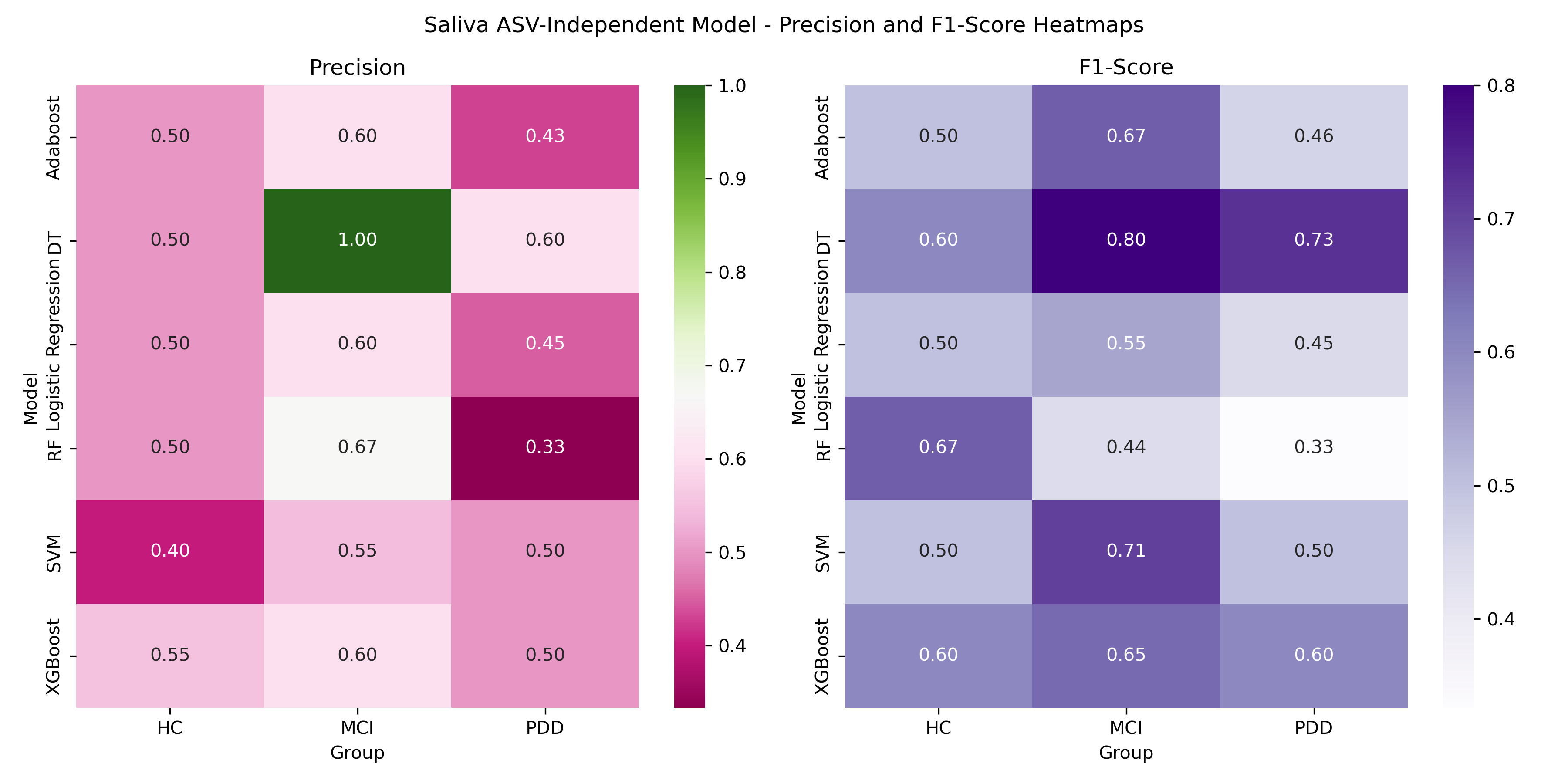

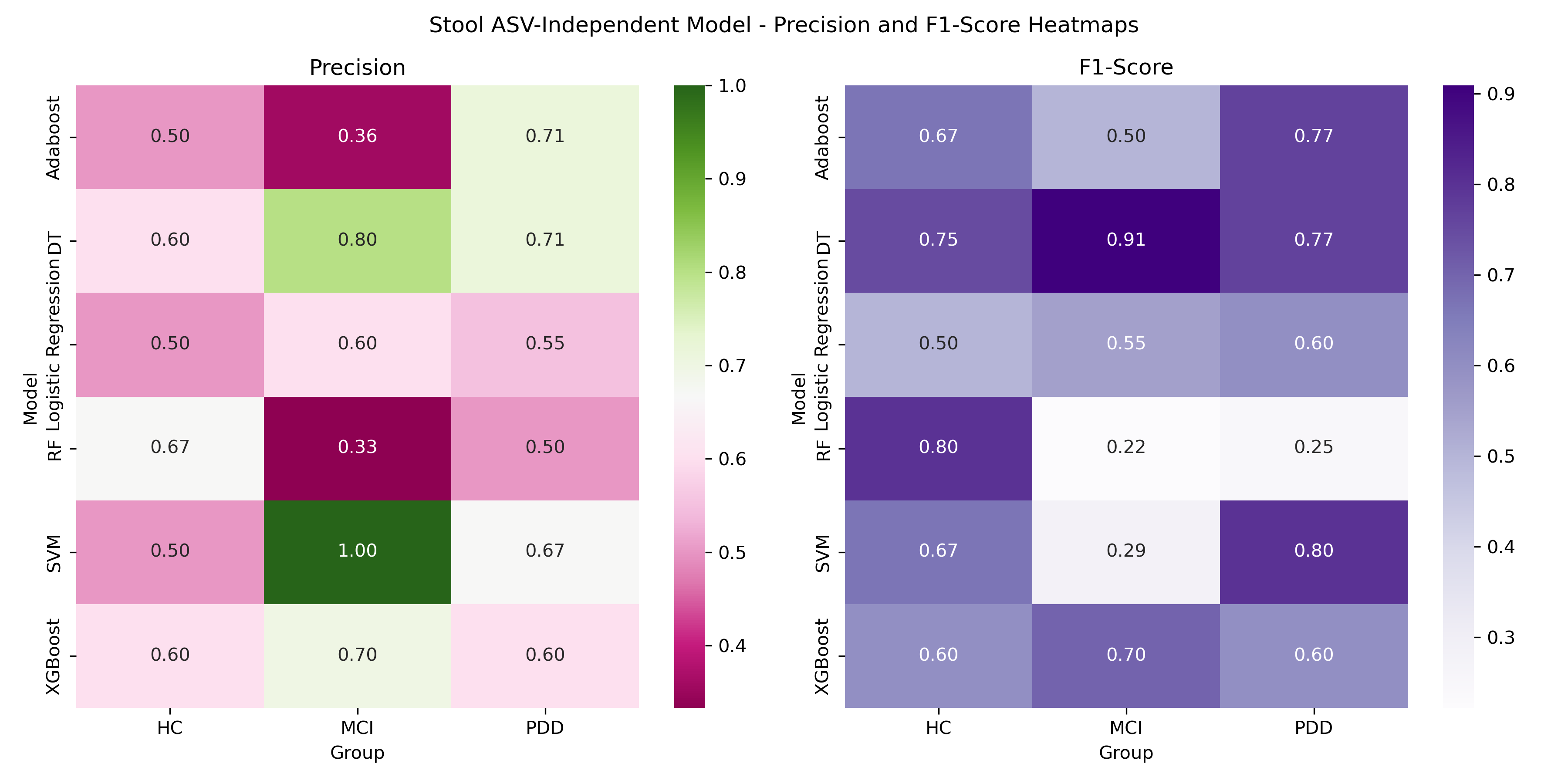

**Table S3**. Topological properties of the selected feature networks between the study groups.

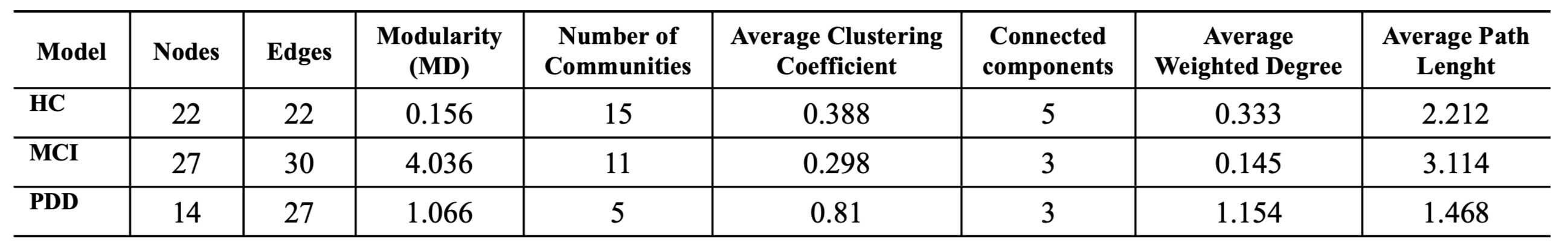

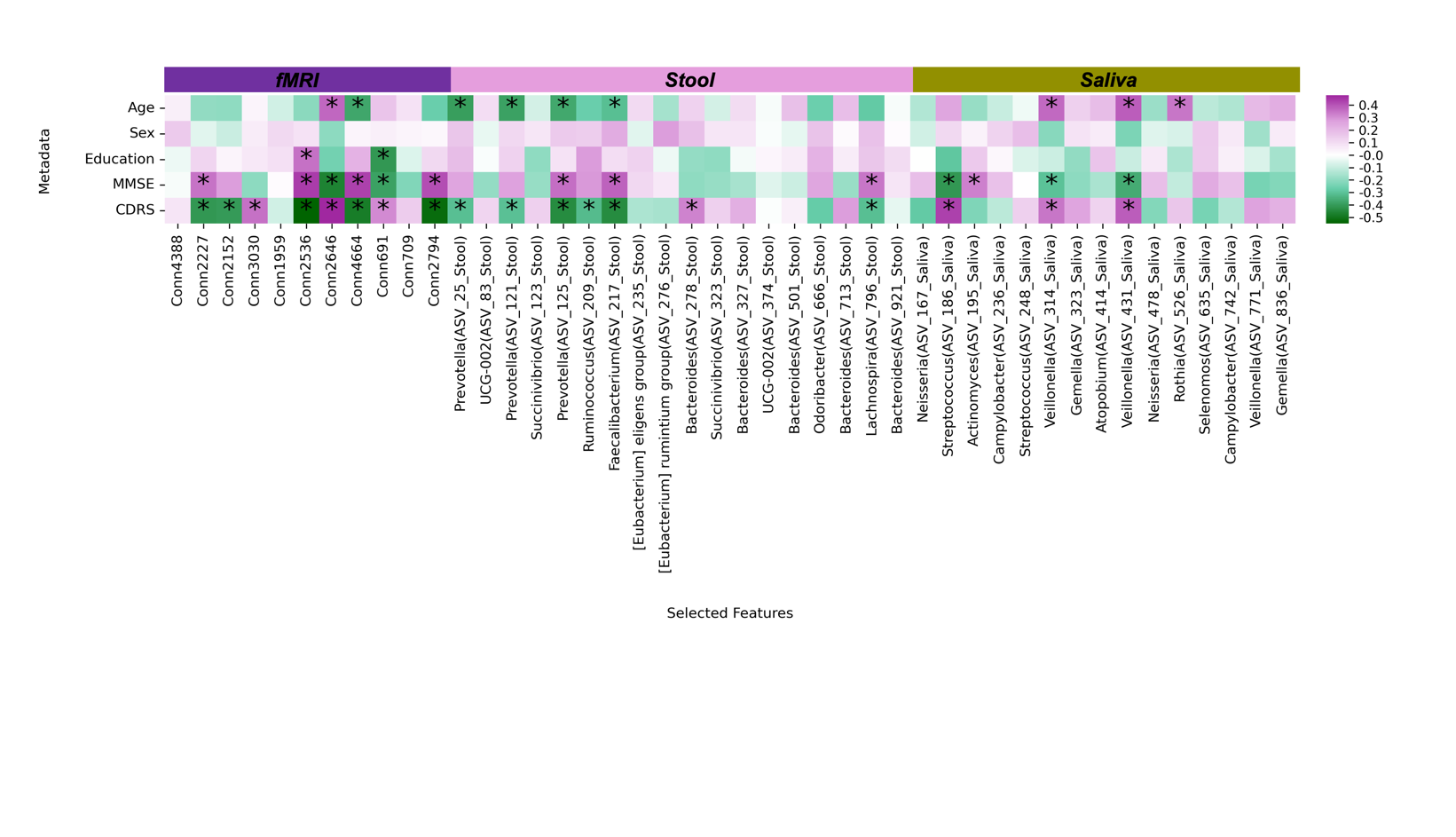
**Figure S3.1.** Associations between metagenomics-selected features, fMRI-selected features and metadata.

a. Heatmap of Spearman correlation analysis between fMRI selected features and metagenomic ASVs and metadata. Correlation coefficients are color-coded, from dark green showing negative correlations to dark purple showing positive correlations. Statistically significant correlations between fMRI-selected features and metagenomic ASVs with |r| ≥ 0.30 and p < 0.05 are asterisked (*) to highlight meaningful associations within the dataset.

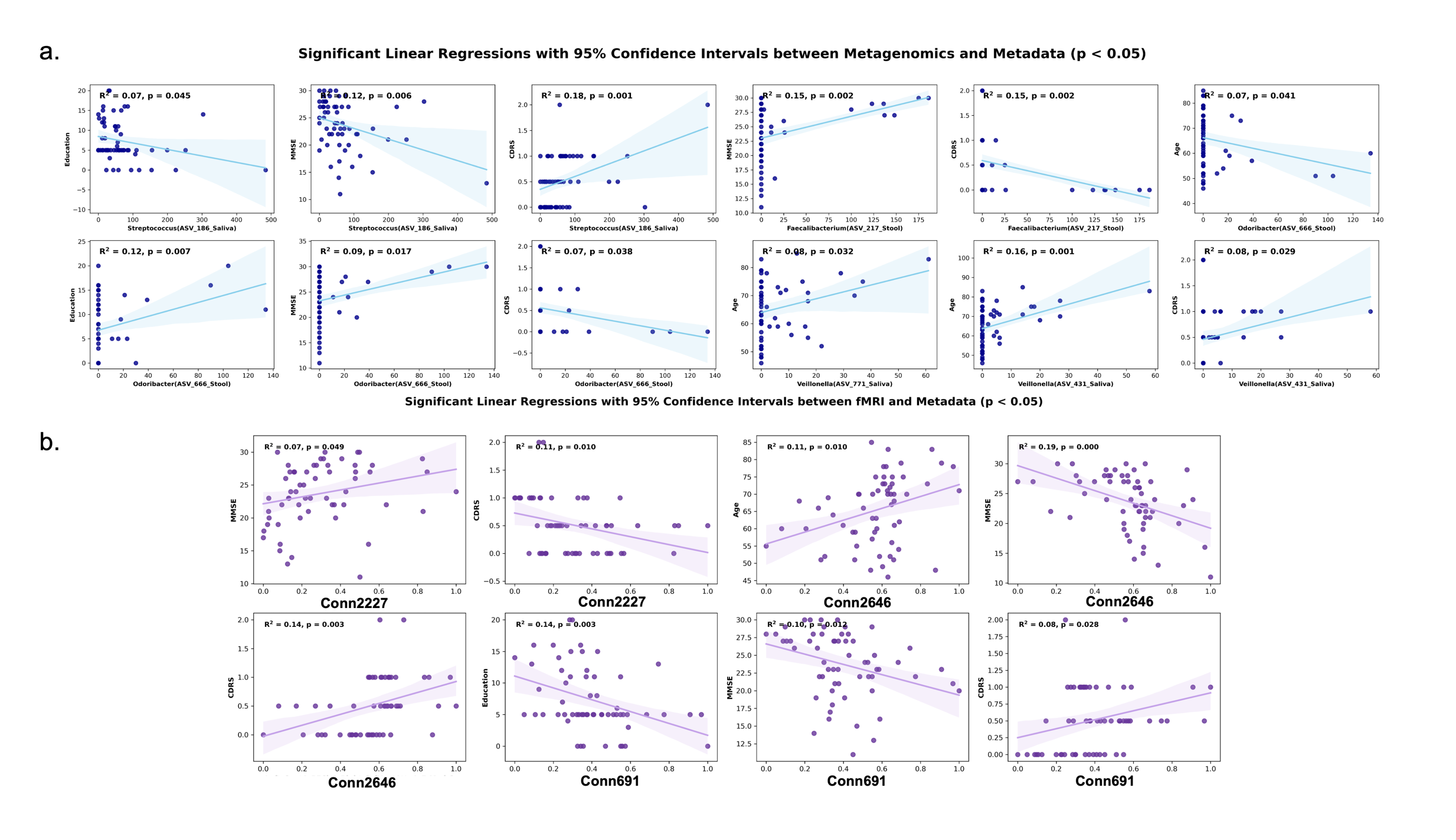
**Figure S3.2.** Significant linear regressions between fMRI-selected features metagenomics-selected ASVs and metadata.

(a) Significant linear regressions between metagenomics-selected features and metadata. (b) Significant linear regressions between fMRI-selected features and metadata. Each scatter plot shows a linear regression line (in blue or lilac) with a 95% confidence interval. R² values and p-values represent the strength and significance of each association, respectively. Associations with a p-value < 0.05 are highlighted to emphasize the statistically significant correlations.
