## Supplementary material for "Integrative Analysis of Neuroimaging and Microbiome Data Predicts Cognitive Decline in Parkinson’s Disease": S1. Supporting Information File. S2. Supporting Information File. Materials and Methods S3. Supplementary Table: S2_Smethods_ML_article.docx

**S1 | MATERIALS AND METHODS**

**S1.1| 16S rRNA Amplicon Sequencing & Analysis**

**S1.1.1| Library generation and sequencing for 16 S rRNA gene amplicon gene sequencing**

Stool and unstimulated saliva samples from the microbiome regions included in our study were collected according to the protocol, as described previously ^2,3^. Fecal and unstimulated saliva samples were performed for microbial DNA extraction using the DNeasy® PowerSoil Kit (QIAGEN GmbH, QIAGEN Strasse 1, 40724 Hilden, Germany).

The saliva samples were centrifuged in 250 ml at 10,000 x g for 5 min, the supernatant was discarded, and the pellet was resuspended with 400 μl of bead beating buffer and transferred to the PowerBead tube. Fecal samples were retrieved from -80 and added to approximately 200 mg PowerBead tubes with a sterile scalpel. Then, the C1 solution (60μl) of the DNeasy® PowerSoil Kit was added.

Samples were homogenized with beads (30 s at level 4, 30 s incubation on ice, and 30 s at level 4) using the Next Advance Bullet Blender. After the bead beating step, the manufacturer’s protocol was followed without any modifications. The V3–V4 regions of the 16 S rRNA gene in all fecal and saliva samples were amplified using universal bacterial primers (F-5′-CCTACGGGNGGCWGCAG-3′ and R-5′-GACTACHVGGGTATCTAATCC-3′). Subsequently, amplicon libraries were prepared following Illumina’s 16 S rRNA metagenomic sequencing library preparation protocol and sequenced using a MiSeq platform and a 2×250 paired-end kit. Amplicon sequencing libraries prepared from a total of 115 gDNA samples were sequenced along with DNA extraction negative control and no-template PCR control for each run ^2,3^.

**S1.1.2 | Processing and analysis of 16 S rRNA gene amplicon sequencing data**

Amplicon sequence data were processed locally using the DADA2 pipeline (14). Quality profiles were filtered according to criteria of (maxN=0) and a minimum length of 50 bp allowing up to 5 expected errors per read. Filtered reads were denoised to obtain amplicon sequence variants (ASVs), followed by pairwise merging of forward and reverse reads to generate full amplicons. Chimeric sequences were removed using and the final ASV table was obtained. Taxonomic assignments were performed by aligning against the SILVA reference database (v.138.1). Sequence processing statistics were monitored at each step and read counts were quality-checked.

After preprocessing the 16S rRNA amplicon sequences using the DADA2 pipeline for quality filtering, we obtained 8980 ASVs for stool samples and 24369 ASVs for saliva samples ^4^. Samples with fewer than 500 counts were removed, and only taxa with an average of more than 5 reads in at least 5% of samples were retained. This yielded a final dataset of 941 ASVs for stool and 884 ASVs for saliva, both with sufficient sequencing depth.

**S1.2 | Neuroimaging Data Pipeline Acquisition & Processing**

**S1.2.1 | Neuroimaging Data Acquisition and Preprocessing**

Functional and structural MRI scans were obtained at Istanbul Medipol University research and training hospital, as described before ^1^. Briefly, a 32-channel head coil was used with standard imaging sequences, following a specific scanning order: (1) localizer, (2) resting-state functional MRI (rs-fMRI), (3) field map, (4) T1-weighted, and (5) T2-weighted structural scans. All scans were performed with participants in the “ON” state, in which they were asked to focus on a point and remain as still as possible. To minimize head movement, spongy pads were used to support the head within the coil. Structural images were acquired in a sagittal plane with a TR/TE of 8.1/3.7 ms, a FOV of 256 × 256 × 190 mm³, and a voxel size of 1 × 1 × 1 mm³ ^1^.

Functional scans included 300 volumes (TR/TE: 2230/30 ms, FA: 77°) with 240 × 240 × 140 mm³ FOV, 3 × 3 × 4 mm³ voxel size, and 35 slices; parameters were modified accordingly after scanner upgrade ^1^.

**S1.2.2 | fMRI Data Preprocessing and Time Series Extraction**

Data preprocessing was performed using the FSL software package. Major steps included the conversion of DICOM into NIfTI format, brain extraction, and then motion correction and smoothing into space using a 5 mm FWHM Gaussian kernel. Functional data underwent high-pass filtering at 150 s and head motion correction using MCFLIRT. Spatial normalization to the MNI152 template was done using both linear and nonlinear registrations with the tools FLIRT and FNIRT, respectively. ICA was performed using the FSL MELODIC tool in order to identify the noise-related components. These latter components were further removed using "fsl_regfilt." In the group analyses, age, brain volume, and LEDD have been included as covariates ^1^.

The identification of ROIs relied on a fine-grained parcellation scheme of the brain into 100 functionally distinct territories: the Schaefer Atlas. After spatially normalizing to the MNI152 template, mean time series data were extracted for each ROI, yielding 100 time series per participant; then, these time series were used in order to calculate the functional connectivity between different brain networks ^1^.
